# A noncanonical glycoprotein H complex enhances cytomegalovirus entry

**DOI:** 10.1101/2024.10.13.617647

**Authors:** Michael J. Norris, Lauren A. Henderson, Mohammed N.A. Siddiquey, Jieyun Yin, Kwangsun Yoo, Simon Brunel, Erica Ollmann Saphire, Chris A. Benedict, Jeremy P. Kamil

## Abstract

Human cytomegalovirus (HCMV) causes severe birth defects, lifelong health complications, and $4 billion in annual costs in the United States alone. A major challenge in vaccine design is the incomplete understanding of the diverse protein complexes the virus uses to infect cells. In *Herpesviridae*, the gH/gL glycoprotein heterodimer is expected to be a basal element of virion cell entry machinery. For HCMV, gH/gL forms a “trimer” with gO and a “pentamer” with UL128, UL130, and UL131A, with each complex binding distinct receptors to enter varied cell types. Here, we reveal a third glycoprotein complex, abundant in HCMV virions, which significantly enhances infection of endothelial cells. In this “3-mer” complex, gH, without gL, associates with UL116 and UL141, an immunoevasin previously known to function in an intracellular role. Cryo-EM reveals the virion-surface 3-mer is structurally unique among *Herpesviridae* gH complexes, with gH-only scaffolding, UL141-mediated dimerization and a heavily glycosylated UL116 cap. Given that antibodies directed at gH and UL141 each can restrict HCMV replication, our work highlights this virion surface complex as a new target for vaccines and antiviral therapies.

## MAIN

Human cytomegalovirus (HCMV) is the prototype β-herpesvirus with ∼80% seroprevalence around the world (Zuhair et al., 2019). Accordingly, HCMV is the primary cause of congenital viral infections, which can result in hearing loss and other neurodevelopmental disabilities. In some underprivileged communities, infection is so abundant that HCMV has been linked to educational achievement gaps among minority groups (Lantos et al., 2018). Among the immunocompromised, HCMV infection is life-threatening (Plotkin and Boppana, 2019). Notably, HCMV vaccine development has been pursued since the 1970’s without success, in large part due to the number of different envelope protein complexes and an incomplete understanding of how they assemble and function in virus entry, cell-to-cell-spread and immune evasion.

In all herpesviruses, the glycoprotein gH plays central roles in cell entry and forms different complexes to mediate entry but is thus far only known to assemble with gL as the core element of receptor binding complexes. One HCMV gH-containing complex is termed “the trimer” (gH/gL/gO), and another is termed “the pentamer”, gH/gL/UL128/UL130/UL131A (Chandramouli et al., 2017; Ciferri et al., 2015; Wang and Shenk, 2005). The trimer and pentamer engage different cell surface receptors (Kabanova et al., 2016; Martinez-Martin et al., 2018) to mediate entry into fibroblasts and non-fibroblasts, respectively. The trimer is stably maintained during serial passage regardless of cell type, but the pentamer is not. Mutations that abolish pentamer expression show a fitness advantage in fibroblasts while pentamer-null viruses exhibit attenuated infectivity in many non-fibroblast cell types (Jiang et al., 2008; Nguyen and Kamil, 2018; Wille et al., 2010). Notably, a third gH complex involving UL116, but not gL, is also present in virions. However, the function, activity, and potential receptors for this gH/UL116 complex remain unknown.

### UL141 is a virion constituent that complexes with gH and UL116

Curiously, although no function is yet known for it, the gH/UL116 interaction is abundant in virions, detected in virions at levels approximating or exceeding those of the gH/gL/gO trimer (Caló et al., 2016; Siddiquey et al., 2021). The abundance of gH and UL116 together suggests that their interaction plays a role of major importance on the viral surface, such as attachment, entry or cell-to-cell spread. However, unlike the trimer and pentamer, the gH/UL116 pair, on its own, fails to bind HCMV-susceptible target cells (Vezzani et al., 2021). Consequently, we wondered if an as-yet-unidentified component might associate with gH and UL116 to mediate viral binding and entry. Supplemental data in a recent mass spectrometry analysis of 170 infected fibroblast cell lines, each stably expressing a single HCMV protein, suggested that gH can associate with a UL141-containing complex in infected cells (Nobre et al., 2019). This broad inventory of the HCMV interactome prompted us to examine whether UL141 might directly interact with gH and UL116 in cells or virions. The HCMV strain TR3 was engineered to encode UL141 fused to a C-terminal FLAG tag. Additionally, we restored a functional UL141 gene to HCMV strain TB40 (TB40^141+^), in the context of a molecular clone that harbors a myc-tagged UL116 (Siddiquey et al., 2021). Parental TB40 (TB40^141-^) is adapted for cell culture, but not for an immunocompetent host, and cannot express UL141 due to frameshift mutation that disrupts the gene (Sinzger et al., 2008; Tomasec et al., 2005). The tagged proteins were readily detected from infected cell lysates (**Fig. 1a-b**). Anti-FLAG immunoprecipitation of UL141 from lysates of TR3 UL141^FLAG^ infected cells pulled down both UL116 and gH, but did not pull down gL. Similarly, anti-myc immunoprecipitation of UL116 from TB40^141+^ virus infected cells pulled down UL141 and gH, but did not pull down gL. These results suggested that gH, UL116, and UL141 together form an envelope protein complex without gL (**Fig. 1b)**. We termed this gH/UL116/UL141 complex the ‘3-mer’.

**Fig. 1.**
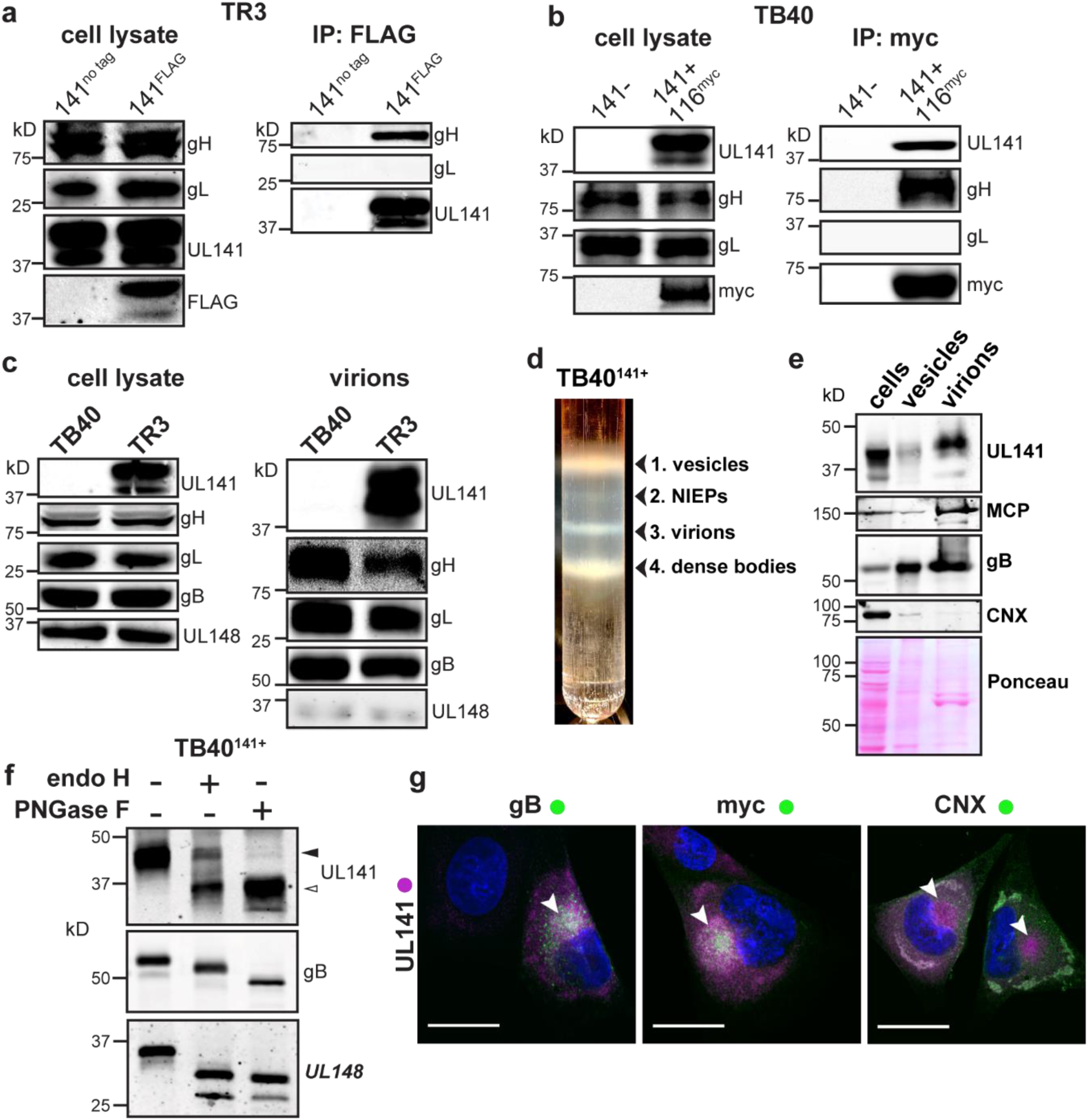
UL141 is incorporated into virions and assembles into a gH/UL116 complex. **a**, The HCMV BAC-derived strain TR3 was modified to introduce a FLAG tag at the C-terminus of UL141 (TR3-141F). Infected fibroblasts were lysed 72 h post infection (hpi), and IP was performed with anti-FLAG. IP eluates were resolved by SDS-PAGE and analyzed by immunoblot with the indicated antibodies. **b**, Fibroblasts were infected with BAC-derived TB40/E (TB40-BAC4) repaired for UL141 expression and engineered to express a myc-tagged UL116 (141+,116myc). Anti-myc IP was carried out at 72 hpi and eluates were analyzed by Western blot. **c**, Infected cell lysates and purified virions from HCMV strains TB40 (141-) and TR3 (141+) were compared for the indicated viral glycoproteins. **d,** TB40-141+ viruses were grown in fibroblasts and concentrated through a 20% sorbitol cushion prior to glycerol-sodium tartrate gradient purification to separate infectious virions (band 3) from cell debris (bands 1 and 4) and non-infectious enveloped particles (NIEPs, band 2). **e**, Western blot analysis of fibroblasts’ whole cell lysates at 6 dpi, vesicles (c, band 1), and virions (c, band 3). Major capsid protein (MCP) identifies the fraction that is enriched for infectious virions. **f**, Lysates of membranes from TB40-141+ infected cells were treated with endoglycosidase H (endoH) or protein N-glycosidase F (PNGase F) and analyzed by Western blot. **g**, Fibroblasts were infected with TB40-141+ at MOI 1 for 3 days prior to staining for UL141 (magenta) and gB, UL116-myc and CNX (all green). The cytoplasmic viral assembly compartment is shown by the white arrowheads, and CNX identifies the ER. Scale bars are 25 μm.

We next tested if the 3-mer could promote cell binding. Given that UL141 can bind CD155 and TNF-related apoptosis inducing ligand (TRAIL) death receptors with nanomolar affinity (Nemčovičová et al., 2013; Tomasec et al., 2005), we hypothesized that UL141 might serve as the receptor-binding component of the gH/UL116/UL141 complex. To begin, we first asked whether UL141 is incorporated into HCMV virions, in addition to its previously known intracellular location where it restricts expression of immune-activating host molecules on the infected cell surface. We readily detected UL141 in virions of strain TR3 but not parental strain TB40 that had not been repaired for the UL141 frameshift (**Fig. 1c**). To further confirm that UL141 is a virion component, HCMV TB40^141+^ virions were isolated by glycerol/sodium-tartrate gradient-purification and detected by anti-capsid antibody (**Fig. 1d-e**). Indeed, UL141 was present in the purified virions, while UL148, an ER-based HCMV glycoprotein which shares a type I transmembrane topology with UL141, was present in only trace quantities in virions (**Fig 1c**). Further, endoglycosidase analysis of cells infected with TB40^141+^ revealed that a substantial portion of UL141 accumulates in an endoglycosidase H (endoH)-resistant form. Endo H cleaves high-mannose, but not complex glycans. Partial resistance of the glycoprotein to endoH-mediated glycan removal is consistent with maturation beyond the ER, where endo-H-resistant, complex glycans decorate canonical virion envelope proteins such as gB (**Fig. 1f**). Third, and consistent with its endo-H resistance, we observed that UL141 localizes to the Golgi-derived cytoplasmic virion assembly compartment (cVAC). The cVAC is where secondary envelopment of progeny herpesvirus virions occurs, and where other structural glycoproteins such as gB and UL116 also localize (**Fig. 1g, Supplemental Data Fig. S1**). Together, these data show that in addition to the role of UL141 in restricting immune activating receptors intracellularly, UL141 also localizes to the cVAC and is incorporated into virions, suggesting an additional role for this immunoevasin, but on the virion surface.

### UL141 facilitates infection of endothelial cells

To address if the 3-mer envelope complex facilitates infection, we tested whether *UL141*-containing TB40^141+^ virions showed enhanced infectivity as compared to the *UL141*-deficient parental TB40^141-^ (**Fig. 2**). We measured absolute infectivity by calculating the ratio of infectious viral particles to viral particles that contain genome but are not infectious, (TCID_50_)/genome (**Fig. 2a-b**). Additionally, we produced virus in both fibroblasts and human umbilical vein endothelial cells (HUVEC) and used this cell-type-specific produced TB40^141+^ and TB40^141-^ to then directly infect both these cell types. Notably, UL141 significantly improves the capacity of HUVEC-derived viruses to subsequently infect endothelial cells but not fibroblasts (**Fig. 2c-e**, **Fig. 3a-b**), which was expected given the trimer is the primary entry factor for fibroblasts. Virions of parental strain TB40^141-^ produced from HUVEC were previously reported to subsequently poorly infect the same cells (Scrivano et al., 2011). A hypothesis at the time was that infected HUVEC produce virions with lower levels of pentamer, which reduces their infectivity for non-fibroblast cells. Our new results suggest that the use of an HCMV strain unable to express UL141 in the past study, and therefore, the absence of the gH/UL116/UL141 3-mer, compromised infectivity for HUVEC.

**Fig. 2.**
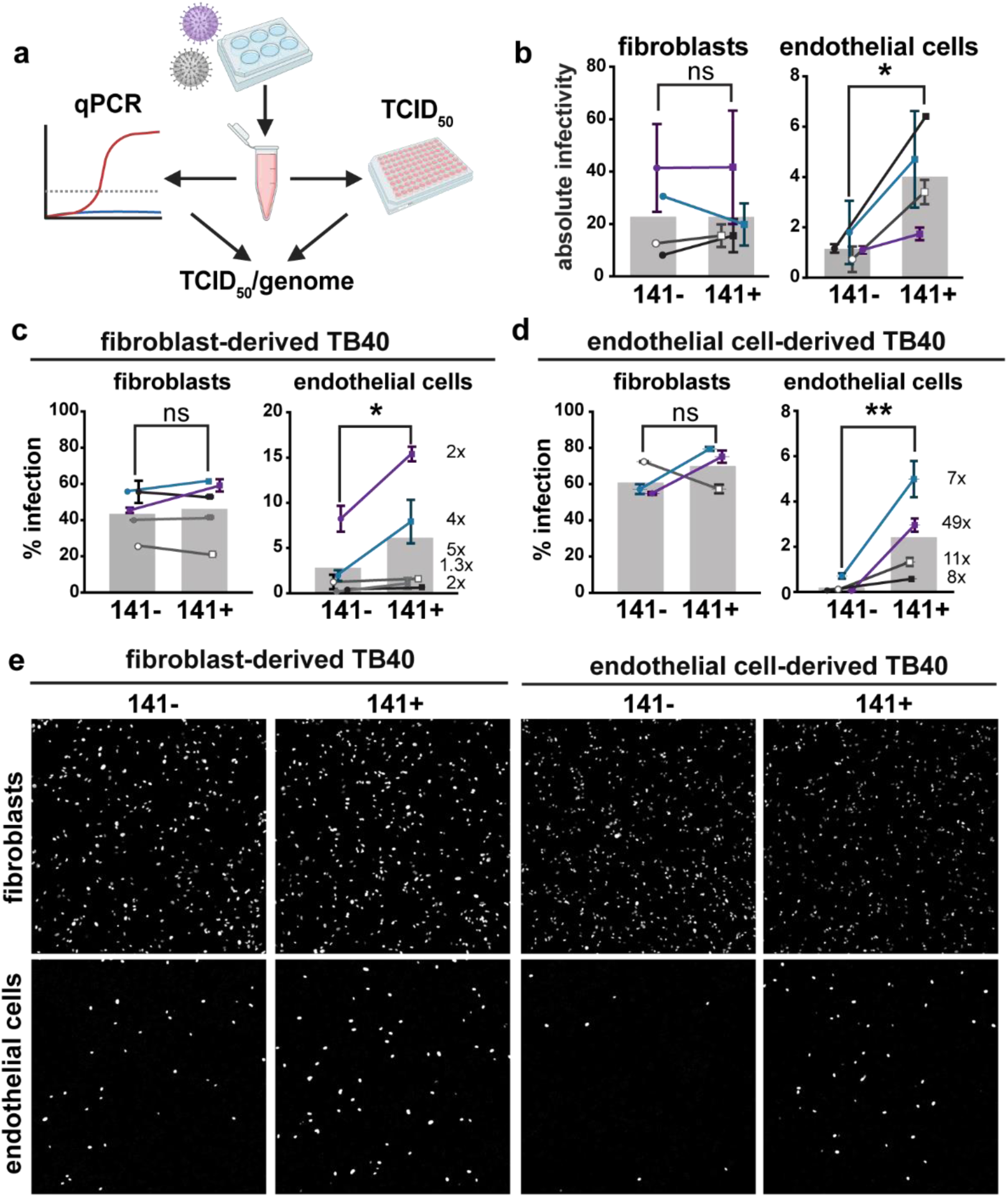
**The 3-mer improves endothelial cell infectivity**. **a,** Schematic of methods used to measure the absolute infectivity of TB40 virions. Viral genomes/mL were calculated from a standard curve with a 10-fold dilution series of TB40 BAC DNA. TCID50 assays were performed on fibroblasts and HUVEC in parallel by staining for IE1 at 3 days post infection, and together were used to calculate TCID50/genome (BioRender.com). **b,** Absolute infectivity shown as TCID50/10^3^ genomes for fibroblasts and endothelial cells, and data represent 4 biological replicates. **c,** Infectivity of fibroblast-derived and **d,** endothelial cell-derived 141+ or 141-HCMV TB40 with fold differences shown as #x for each biological replicate. Fibroblasts and endothelial cells were infected for 24h in parallel with 50 or 100 genome equivalents/cell, respectively, stained for IE1 and counterstained with Hoechst 33342 to calculate the percentage of infected cells. **e,** Representative images of c and d. Statistical significance was determined by a paired t-test, with each point representing a biological replicate consisting of 2-3 technical replicates each. Error bars represent ± SEM for each biological replicate. Shaded bars are the mean % infection for all biological replicates.

**Figure 3.**
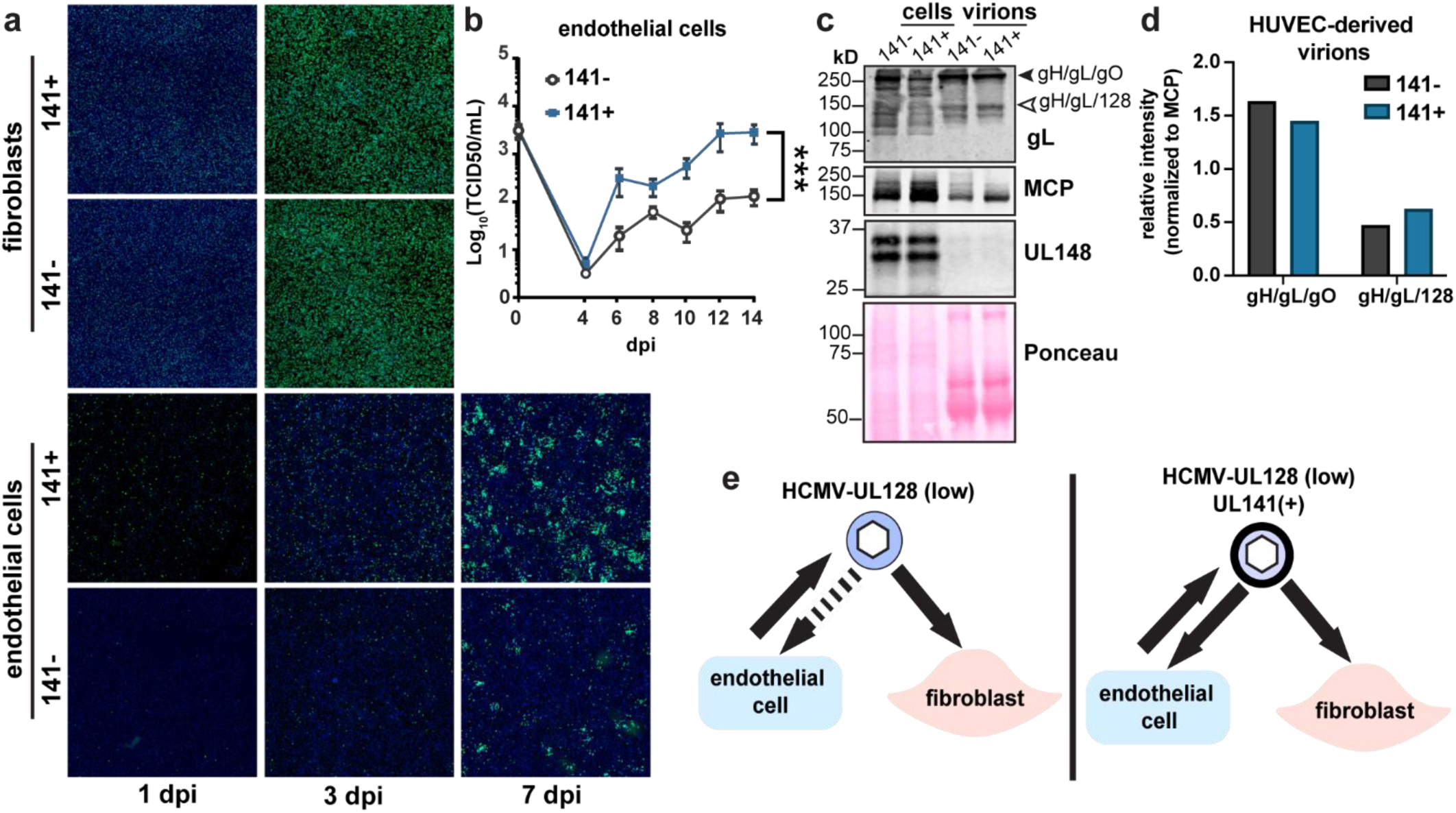
UL141 promotes endothelial cell tropism independently of the pentamer complex. a, Representative images of UL141-dependent spread in endothelial cells. Fibroblasts and endothelial cells were infected with 50 or 100 genome equivalents/cell, respectively, of 141-or 141+ TB40 viruses produced by fibroblasts. Cells were stained for IE1 (green) and Hoechst (blue) at the indicated days post-infection to monitor viral spread. **b,** Low MOI (0.01 TCID50) multicycle viral growth kinetics in HUVEC that were infected with 141-or 141+ viruses up to 14 dpi. This graph is a compilation of 3 biological replicates. Lognormal data were logarithmically transformed to fit a Gaussian distribution prior to calculating statistical significance via 2-way ANOVA for 12 dpi data points. **c**, Non-reducing SDS-PAGE of HUVEC cell lysates and HUVEC-derived virions. Cells were infected with 141– and 141+ viruses to measure virion incorporation of known HCMV entry complexes, trimer (gH/gL/gO) and pentamer (gH/gL/128). HUVEC-derived virions were concentrated through a 20% sorbitol cushion at 12 dpi and 14 dpi prior to lysis. Whole cell lysates were collected at 14 dpi. Lysates were immunoblotted for gL to identify covalently linked entry complexes, major capsid protein (MCP) to measure virion abundance, and UL148 to assess the purity of the virion preparations. **d**, Quantification of band intensities for gH/gL/gO and gH/gL/128 abundance in HUVEC-derived virions from **c**. Band intensities of 141– and 141+ virions were normalized to MCP. **e**, Graphical summary illustrating the role of UL141 as an endotheliotropic factor that restores the ability of TB40 virions to subsequently infect endothelial cells.

Importantly, the pentamer complex is similarly incorporated into UL141– and UL141+ virions produced in endothelial cells, suggesting that UL141 enhances endothelial cell tropism in a pentamer-independent manner (**Fig. 3c-e**). To investigate this further in another cell type that utilizes the pentamer for efficient viral internalization, we infected epithelial cells with the pentamer-null HCMV AD169 strain. In addition to carrying a frameshift mutation in UL131A that prevents expression of the pentamer, AD169 lacks most of the UL/b’ region encoding *UL141* and ∼15 other genes (**Fig. 4a**). We restored *UL141* to this strain and found that epithelial cells infected with UL141-restored AD169 (AD169^141+^) produce larger foci in comparison to those infected with AD169^141-^ (**Fig. 4b-c, Supplemental Data Fig. S2**). These data indicate that UL141 augments infection in the presence or absence of the pentamer complex.

**Fig. 4.**
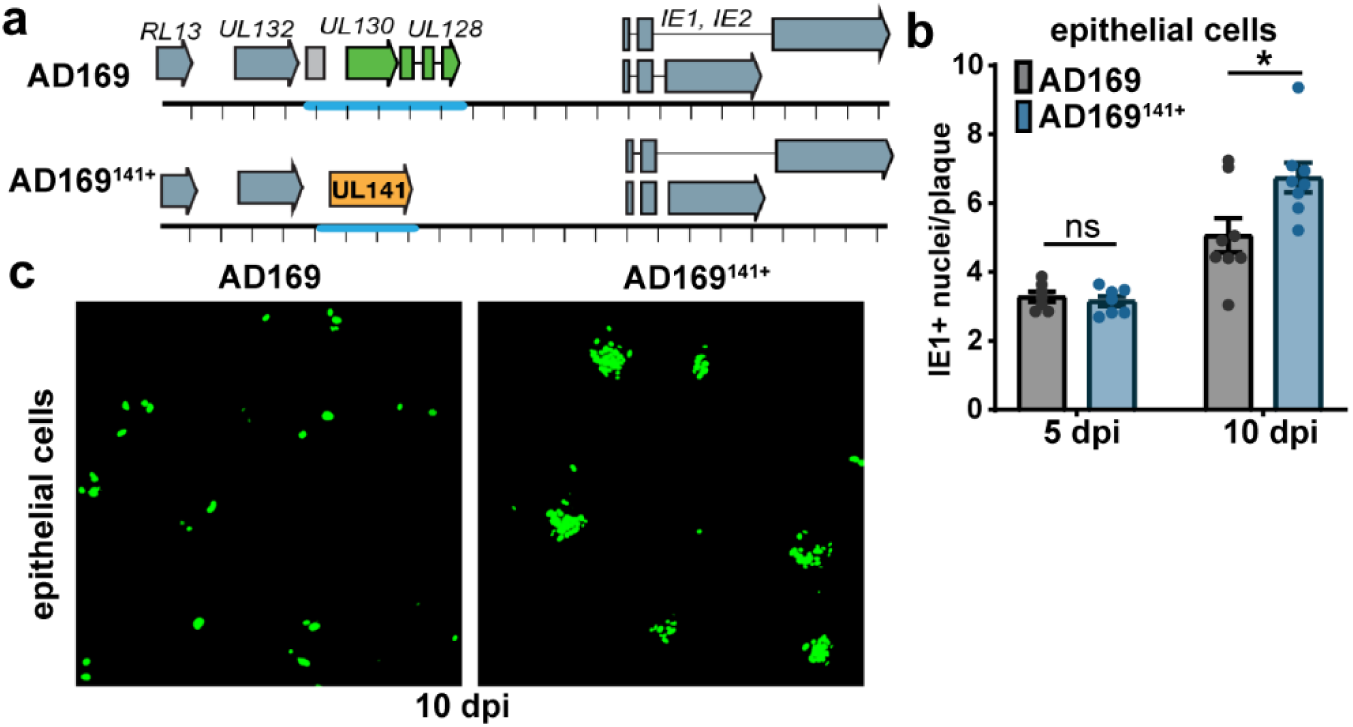
UL141 enhances the infectivity of the pentamer-null AD169 strain. a, Schematic of the pentamer-null AD169 viruses used in the following experiments. **b**, ARPE-19 were infected with AD169 or UL141-restored AD169 (AD169^141^) at MOI 0.1. Cells were stained for IE1 at 5 and 10 dpi to measure the size of foci, or plaques in IE1+ nuclei per plaque. Each point in the bar graph represents a biological replicate (N=7-8). Data was logarithmically transformed to fit a Gaussian distribution prior to measuring statistical significance via Welch’s t test. **c**, Representative image of AD169 versus AD169^141+^ plaques in ARPE-19 cells at 10 dpi. Cells were stained for IE1 (green).

To exclude the possibility that UL141 facilitates entry independently of gH/UL116, we generated UL116-deficient (Δ116) TB40 viruses that were either *UL141-* or *UL141+*. In contrast to *UL116*+ virus, HCMVΔ116 showed no enhancement of infectivity when restoring *UL141*. We also saw that in the absence of UL116, UL141 is poorly incorporated into the virion, even though it is still trafficked to the cVAC (**Fig. 5a-d**). Together, these results suggest that assembly of UL116 with UL141 in the 3-mer facilitates UL141-driven endothelial cell tropism.

**Fig. 5.**
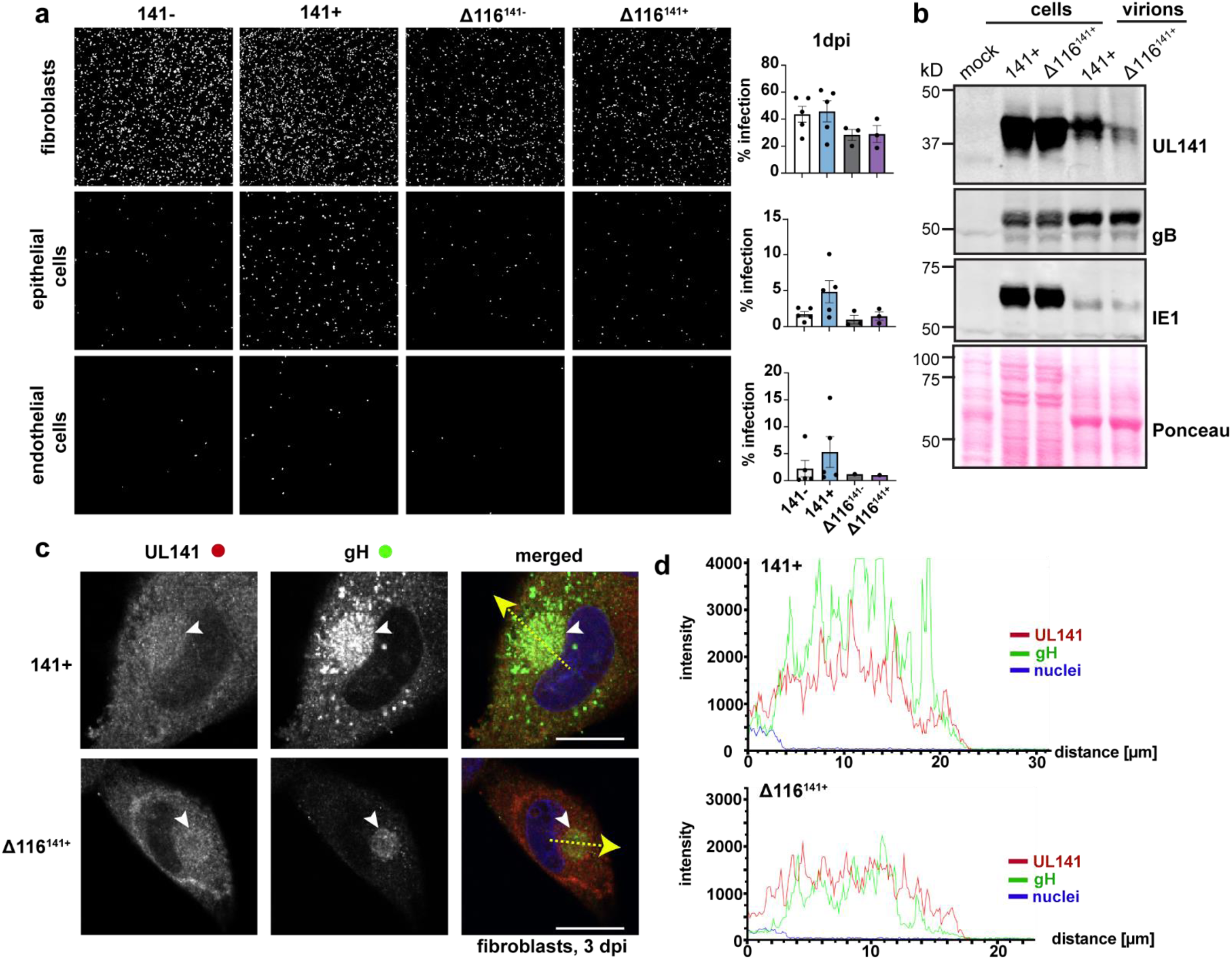
UL116 is required for UL141-dependent entry and virion incorporation of the 3-mer. a, Representative images of fibroblasts, epithelial cells, and endothelial cells following infection with UL116-sufficient versus UL116-deficient (Δ116) TB40 that encode UL141 or not (141– and Δ116^141-^; 141+ or Δ116^141+^). Fibroblasts and epithelial cells (ARPE-19) were infected with 50 genome equivalents/cell, while endothelial cells (HUVEC) were infected with 100 genome equivalents/cell. The cells were stained for IE1 and counterstained with Hoechst at 1 dpi to measure the percentage of infected cells in each condition. Each point represents a biological replicate. **b**, Western blot analysis of whole cell and crude virion lysates from 141+ or Δ116^141+^ infected fibroblasts. **c,** Immunofluorescence of the cytoplasmic viral assembly compartment (cVAC, white arrowheads) in fibroblasts infected with 141+ or Δ116^141+^ at 3 dpi (MOI 1). Scale bars represent 20 μm. **d**, Intensity profiles of UL141 (red) and gH (green) throughout the cVAC. The regions of interest used to measure the intensity profiles for each condition are represented by the yellow dashed arrows in **c.**

### Structure of the HCMV gH/UL116/UL141 3-mer

To determine the high-resolution structure of the 3-mer, we recombinantly co-expressed the soluble ectodomains of gH, UL141, and UL116 in Freestyle 293F cells and purified the complex by affinity chromatography followed by size exclusion (**Fig. 6a-c**). The resulting heterotrimer is non-covalently linked and dissociates into single subunits in SDS-PAGE under non-reducing conditions (**Fig. 6b and c**). We used single-particle cryo-electron microscopy (cryo-EM) to resolve the structure of the gH/UL116/UL141 complex (**Fig. 6d**; **Fig. 7**; **Fig. 8**; **Fig 9**). Two-dimensional (2D) class averaging revealed a high degree of detail, including secondary structure features, and illustrated a single particle population in the overall shape of a twisted H, in which each gH/UL116/UL141 heterotrimeric complex forms a dimer with a second gH/UL116/UL141 heterotrimer (**Fig. 6d**). The structure of the dimeric gH/UL116/UL141 heterotrimeric complex was determined to a resolution of ∼3.5 Å (**Fig. 7**; **Fig. 8**; **Fig. 9**), allowing us to build a structural model of the majority of the residues of the gH, UL141, and the previously unknown UL116 subunits. Symmetry expansion and focused 3D reconstructions on gH-UL116 enabled further structural modeling in this region, including the identification of several glycans that were unresolved in the dimer map (**Fig. 7b & c inset; Fig 8a-d**). The N-terminal domain of UL116 (residues 1-202) could not be resolved in our cryo-EM structure, despite its presence in the protein construct and sample. This is likely due to the inherent flexibility of this region and the predicted extensive glycosylation. Specifically, there are fourteen predicted consensus sites for N-linked glycosylation and over 70 potential sites for O-linked glycosylation, which may contribute to the observed structural disorder (Caló et al., 2016).

**Fig. 6.**
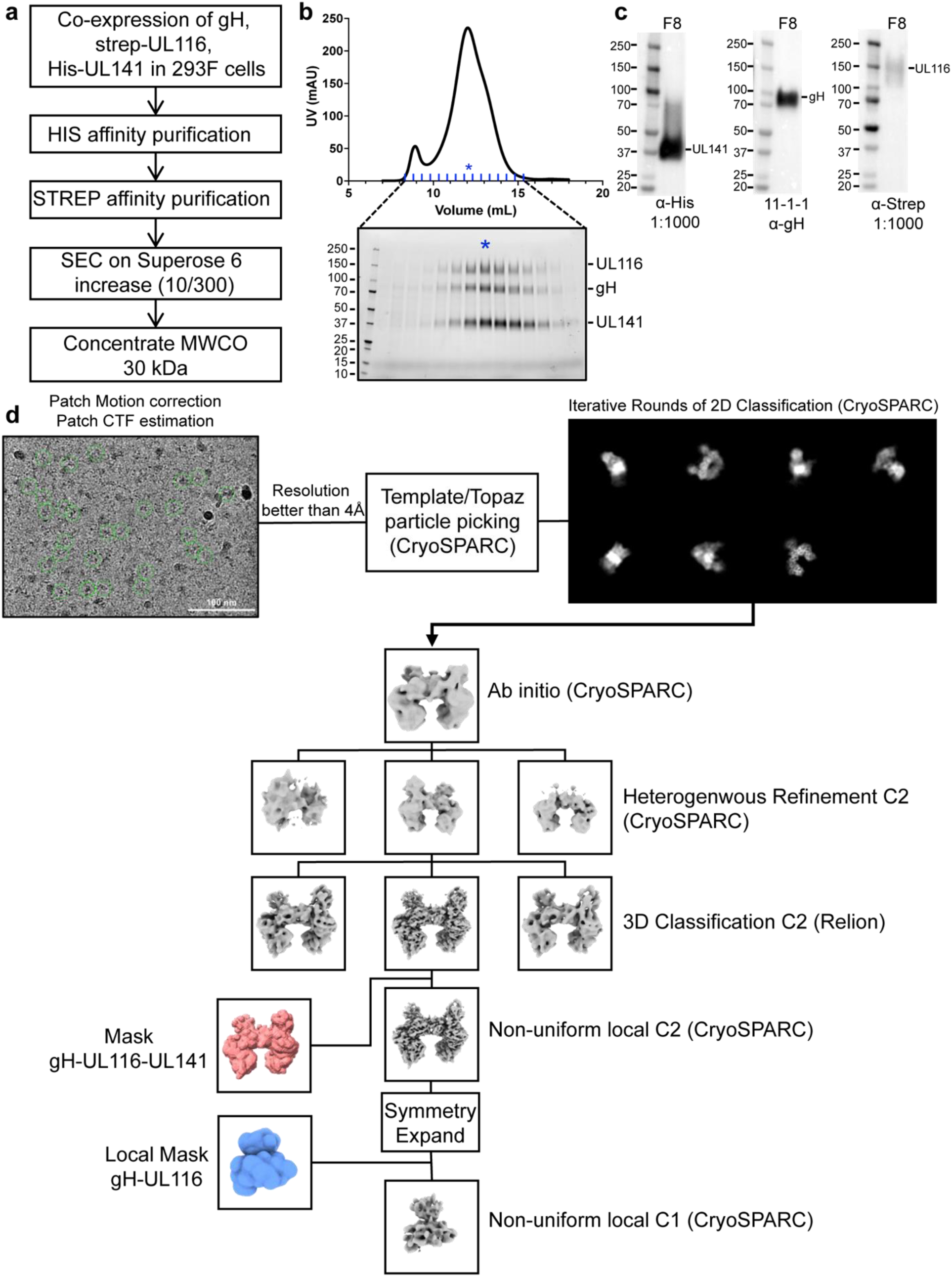
Purification and cryo-electron microscopy processing of the HCMV gH/UL116/UL141 3-mer. a, Schematic representation of the expression and purification process for the HCMV gH/UL116/UL141 complex. **b,** Size exclusion chromatography profile of the gH/UL116/UL141 complex. Fractions were analyzed by SDS-PAGE under non-reducing conditions, with the fraction indicated by blue asterisks used for cryo-EM studies. **c,** Western blot analysis of fraction 8, probed with anti-His, anti-gH, and anti-strep antibodies to detect UL141, gH, and UL116, respectively. **d,** Overview of the representative cryo-EM data processing workflow for the gH/UL116/UL141 complex.

**Fig. 7.**
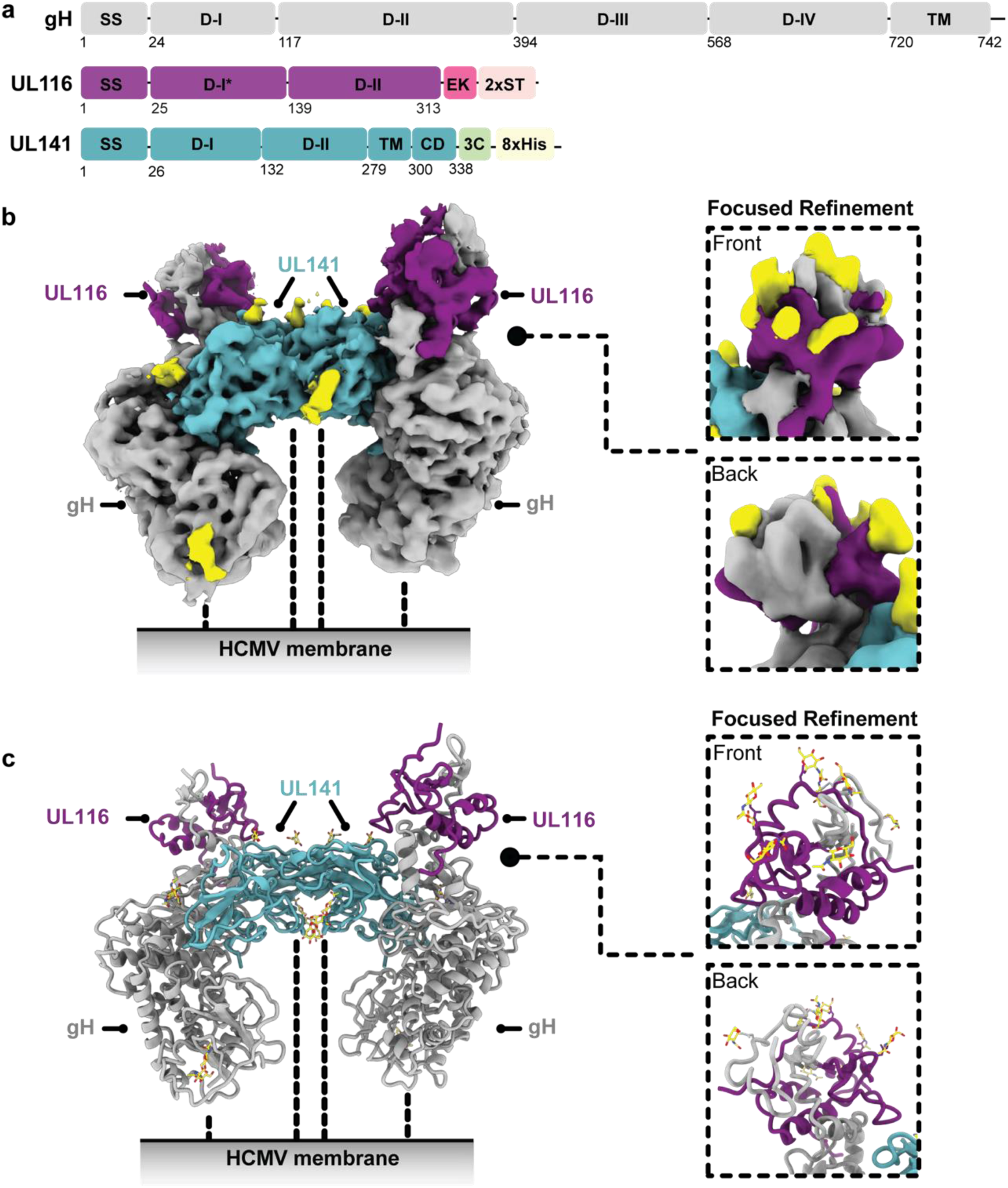
**Cryo-EM structure of HCMV gH/UL116/UL141 3-mer**. **a,** Schematic representation of the domain organization of HCMV gH, UL116, and UL141. **b**, Cryo-EM map of the HCMV gH/UL116/UL141 3-mer complex ectodomain, with gH shown in grey, UL116 in purple, and UL141 in teal. The dashed lines indicate the hypothetical locations of the protein stalks. The inset displays front and back views of a symmetry-expanded focused local refinement around the gH-UL116 interaction. Resolved locations of N-linked glycans from focused refinements are highlighted in yellow. **c**, Ribbon diagram of the gH/UL116/UL141 3-mer. The inset presents front and back views of the ribbon diagram focusing on the gH-UL116 interaction. Resolved N-linked glycans from focused refinements are also highlighted in yellow.

**Fig. 8.**
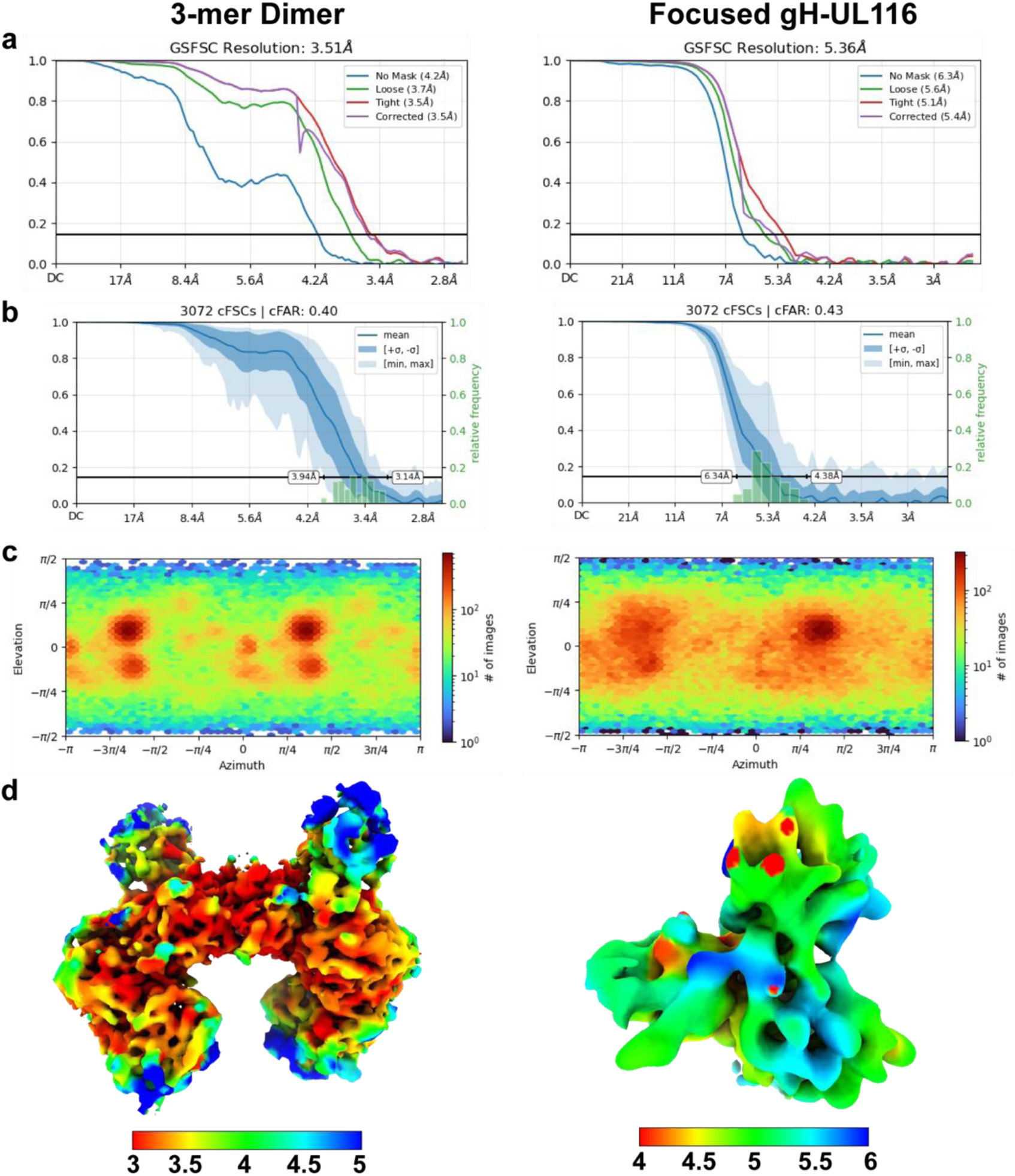
Cryo-EM structure validation. a, Gold-standard Fourier shell correlation (FSC) curves for the refinements of the HCMV gH/UL116/UL141 dimer (left) and the symmetry-expanded focused local refinement of the gH-UL116 interface (right). **b,** Conical FSC (cFSC) analysis of the half maps. The blue cFSC summary plot displays the mean, minimum, maximum, and standard deviation of correlations at each spatial frequency. The green histogram shows the distribution of 0.143 threshold crossings, corresponding to the spread of resolution values across different directions. **c,** Euler angle distribution plot of the particles used in the final 3D reconstructions, demonstrating complete coverage of projections as generated in CryoSPARC. **d,** Final reconstructions filtered and colored by local resolution, as estimated in CryoSPARC.

**Fig. 9.**
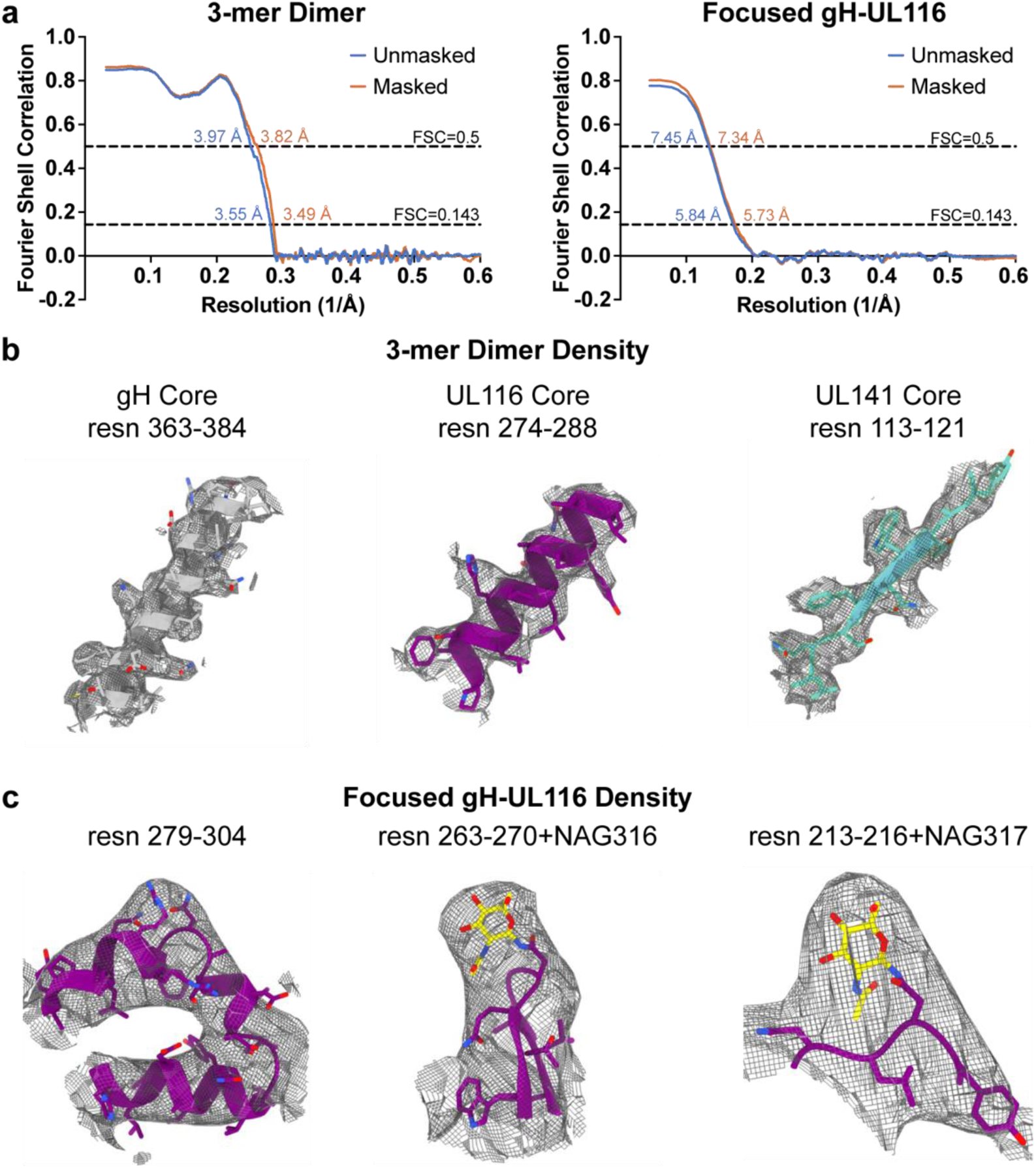
Cryo-EM structure validation and model quality assessment. a, Map versus model FSC curves calculated with and without masking, using the Phenix package. Curves are shown for the HCMV gH/UL116/UL141 dimer (left) and the symmetry-expanded focused local refinement of the gH-UL116 interface (right). **b,** Stereo views of cryo-EM density maps for fragments of gH (left), UL116 (middle), and UL141 (right) from the 3-mer dimer, demonstrating the quality of the density. The cryo-EM density is displayed as a mesh. **c,** Stereo views of cryo-EM density maps for a fragment of gH and UL116 from the symmetry-expanded focused refinement of the gH-UL116 interface, illustrating the quality of the cryo-EM density. The density is shown as a mesh.

The HCMV gH/UL116/UL141 3-mer overall is ∼130 Å in length and 75 Å in width (**Fig. 7**). Unlike the trimer and pentamer, for which the component molecules are organized linearly, like train cars, the 3-mer components do not interact in a linear arrangement. Instead, gH scaffolds both UL116 and UL141 subunits, with gH at the center, UL116 at the top of gH and UL141 at the side of gH (**Fig. 7**). The gH/UL116/UL141 complex exhibits a predominantly negative charge at the viral membrane proximal region (gH) and a small positively charged patch at the distal region (UL116) (**Fig. 10a**). The gH DII domain and UL141 together form a negatively charged cleft along one side of the complex, while the opposite side remains approximately neutral (**Fig. 10a**). This uneven distribution of surface charge could play a crucial role in how the complex engages with its receptor. Moreover, the twelve observed N-linked glycosylation sites are distributed primarily across the top of the gH/UL116/UL141 3-mer complex (**Fig. 10b**). We could confidently model four of the predicted six sites on gH, all three predicted sites on UL141, and five of the fourteen predicted sites on UL116. There is only a single glycan site at the membrane proximal region of the complex, the rest are restricted to the upper portion of the complex. Due to the inherent low resolution of the focused gH-UL116 map, we were unable to confidently model any O-linked glycan sites on UL116. Nonetheless, the observed uneven distribution of glycans, concentrated mainly at the upper portion of the complex, suggests an evolutionary advantage, likely in modulating interactions with cell surface lectins (Feng et al., 2022; Koehler et al., 2020; Li et al., 2021), and may enhance the complex’s ability to evade immune detection by forming a protective glycan shield.

**Fig. 10.**
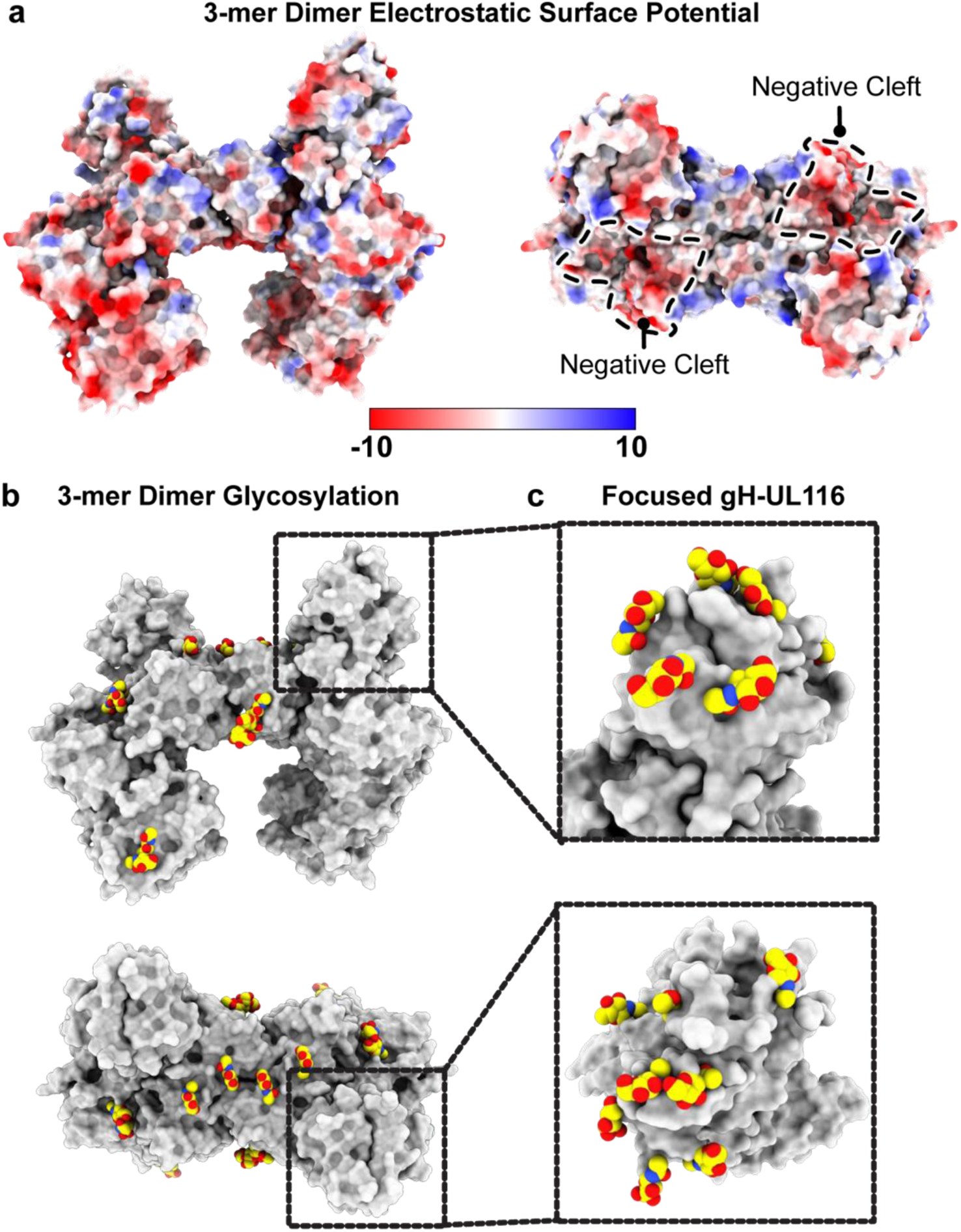
Electrostatic surface potential and glycosylation of the HCMV gH/UL116/UL141 3-mer. a, Electrostatic surface potential of the HCMV 3-mer displayed on a space-filling model, with positively charged regions in blue and negatively charged regions in red. The negatively charged cleft is outlined. Electrostatic potential maps were generated using the PDB2PQR and APBS software. **b,** Side and top views of the glycosylation site distribution on the HCMV gH/UL116/UL141 3-mer. **<u>c,</u>**Inset showing the glycosylation site distribution at the gH-UL116 interaction site, as resolved in the symmetry-expanded focused refinement of the gH-UL116 interface.

The gH component, shared among all three HCMV gH complexes, contains four domains (DI-DIV) (**Fig. 11a**). Each domain, as an individual building block, retains a similar fold in the three complexes (**Fig. 11b**), and all extend linearly away from the membrane proximal face. However, the distal DI is positioned differently in the 3-mer compared to the trimer and pentamer. A significant rotational change is observed in DI, transforming the gH subunit from a straight rod in the trimer and pentamer structures into a crescent shape in the 3-mer structure (**Fig. 11a**). This alteration results in the two beta sheets in gH DI shifting from a horizontal orientation to a vertical orientation and angling of DII in the 3-mer. This adjustment accommodates binding of gH to UL116 to form the 3-mer, as opposed to binding gL to form the trimer and pentamer. Another structural difference is that DI of gH co-folds with UL116 at the membrane distal region of the 3-mer, but co-folds with gL in both the trimer and pentamer. The co-folding suggests that the distinct complexes form prior to their expression on the virion or cell surface.

**Fig. 11.**
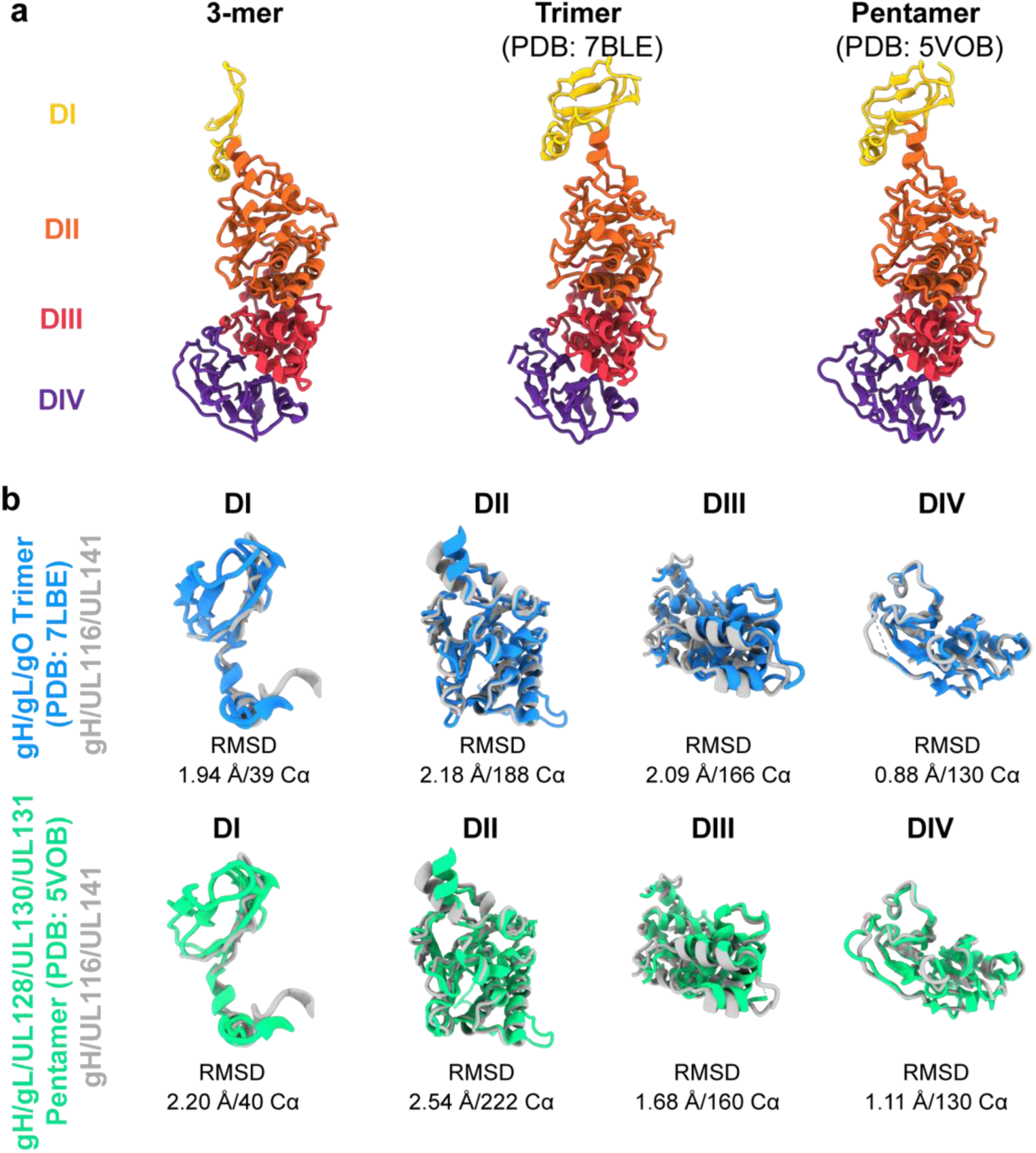
Structural comparison of gH from the HCMV 3-mer, trimer, and pentamer. a, Structural representation and domain organization of gH in the HCMV 3-mer (left), trimer (middle), and pentamer (right). The gH domains I–IV are colored yellow, orange, red, and purple, respectively. In the 3-mer, the gH DI domain undergoes a significant rotational shift relative to the trimer and pentamer, transforming the gH subunit from a straight rod in the trimer and pentamer to a crescent shape in the 3-mer structure. **b,** Structural alignment of individual gH domains comparing the 3-mer with the trimer (top) and the pentamer (bottom). The structures were aligned using the indicated number of Cα atoms from the respective PDB files, and the alignment was quantified by the indicated r.m.s.d. values.

Although it was previously known that gH could interact with UL116 independently of gL (Caló et al., 2016), the specific organization of these proteins relative to each other and the details of their interacting surface contacts, and the structure of UL116, itself, remained entirely unknown. Here, we show the first high-resolution description of UL116 and reveal that it forms a five-stranded beta-sheet together with the N-terminal residues of gH DI. Additionally, a C-terminal alpha-helix of UL116 interacts extensively with the top of gH DII, together forming a capping structure around gH and covering approximately 2492 Å^2^ of its surface (**Fig. 12a**). The binding footprint of UL116 on gH largely overlaps with that of gL in the trimer and pentamer (roughly 80% of the UL116 binding footprint), indicating that the presence of UL116 would preclude simultaneous binding of gL (**Fig. 12b**). Consequently, gH has evolved to recognize both UL116 and gL in distinct complexes, achieved via a structural switch centered around DI. This structural switch allows the HCMV viral surface to be loaded with multiple distinct glycoprotein complexes, expanding cellular tropism and perhaps also confounding immune responses.

**Fig. 12.**
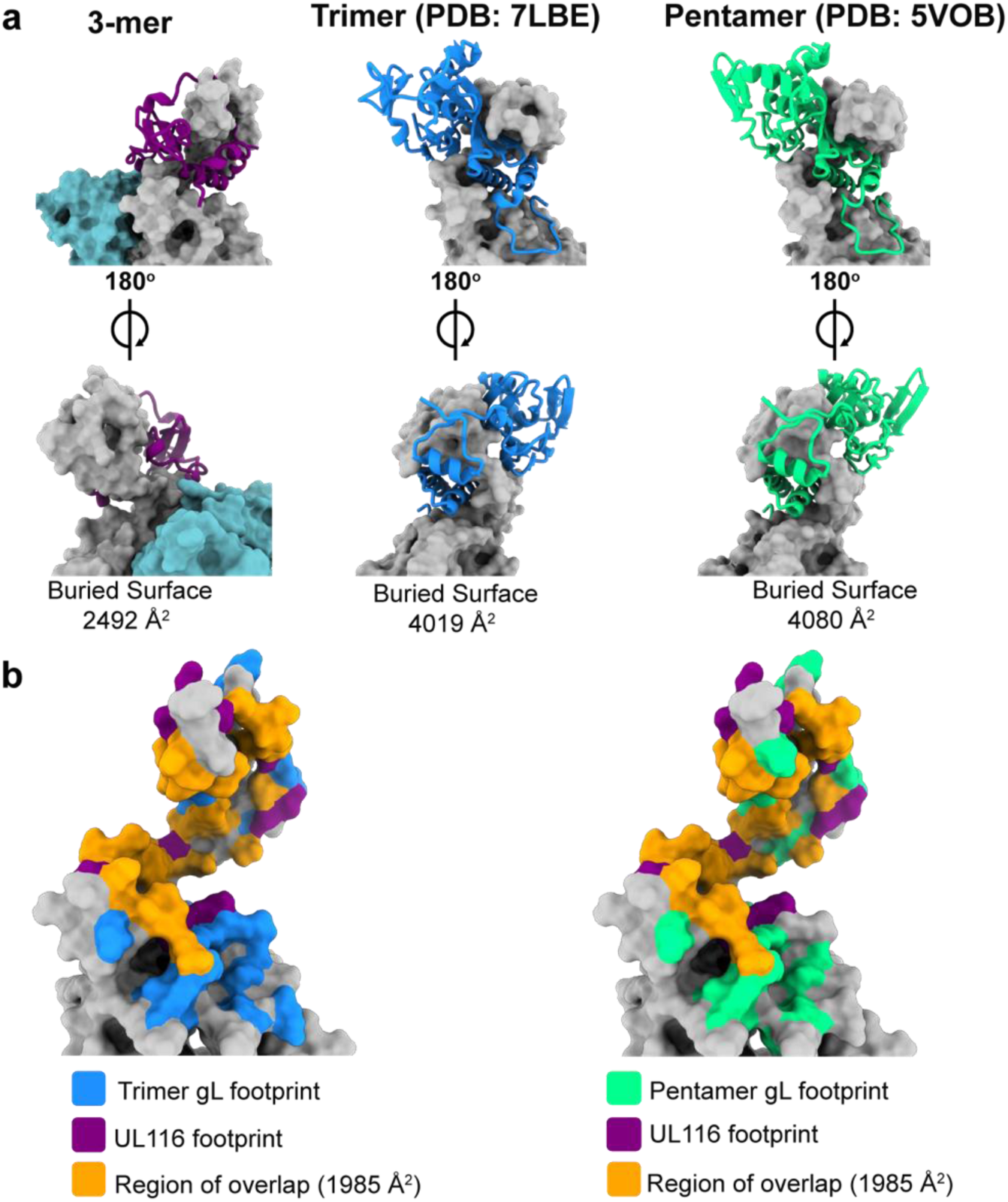
UL116 and gL share similar binding sites on gH. a, Front (top) and back (bottom) views of UL116 (left), gL in the trimer (middle), and gL in the pentamer (right) bound to gH. UL116 and gL are depicted as ribbon diagrams, while gH is shown as a surface model. The calculated buried surface area of each interaction pair is indicated below each structure. **b,** Comparison of the UL116 binding footprint on gH with that of gL from the trimer (left) and the pentamer (right). The UL116 binding footprint is highlighted in purple, gL from the trimer in blue, and gL from the pentamer in green, with the overlapping region shown in orange. The buried surface area of the overlapping region is indicated.

The soluble HCMV UL141 ectodomain forms a dimer in isolation and when complexed with one of its cellular ligands, TRAIL-R2 (PDB: 4JM0 and 4I9X, respectively). Consistent with these findings, our cryo-EM analysis of the 3-mer reveals that the UL141 ectodomain similarly forms a UL141-UL141 dimer, like the crossbar of an “H”, that brings together two independent gH/UL116/UL141 complexes. The UL141 dimeric interaction is non-covalent, head-to-tail, and has a C alpha RMSD of only 0.78 Å when compared with unbound or TRAIL-R2-bound UL141 dimers (**Fig. 13a**). Within the UL141 structure of the 3-mer, however, several regions previously disordered and absent from both the unbound and TRAIL-R2-bound UL141 are well-ordered, including the C-terminal loops 168-174, 199-207, and 217-226. Both the 168-174 and 199-207 loops are part of the extensive gH-interacting surface in the 3-mer but were solvent exposed and mobile when unbound or in complex with TRAIL-R2. At this interaction site, gH and UL141 bury 1985 Å of molecular surface (∼15% of the entire UL141 surface) (**Fig. 13b**). These loops interact with gH DII, while loop 168-174 additionally contacts DIV, and loop 199-207 DIII. The footprint of gH on UL141 significantly overlaps with that of TRAIL-R2 on UL141 (roughly 25% of the TRAIL-R2 footprint) (**Fig. 13c**), indicating that the 3-mer may need to disassemble or undergo a structural change for UL141 to bind TRAIL receptors on the cell surface. The large footprint of UL141 on gH partially obscures the binding site of the gH-neutralizing antibody 13H11 (Kschonsak et al., 2022, 2021), and likely other neutralizing anti-gH antibodies. Antibodies like 13H11 would thus only likely neutralize HCMV entry mediated by the trimer and pentamer, but not entry or spread mediated by the 3-mer.

**Fig. 13.**
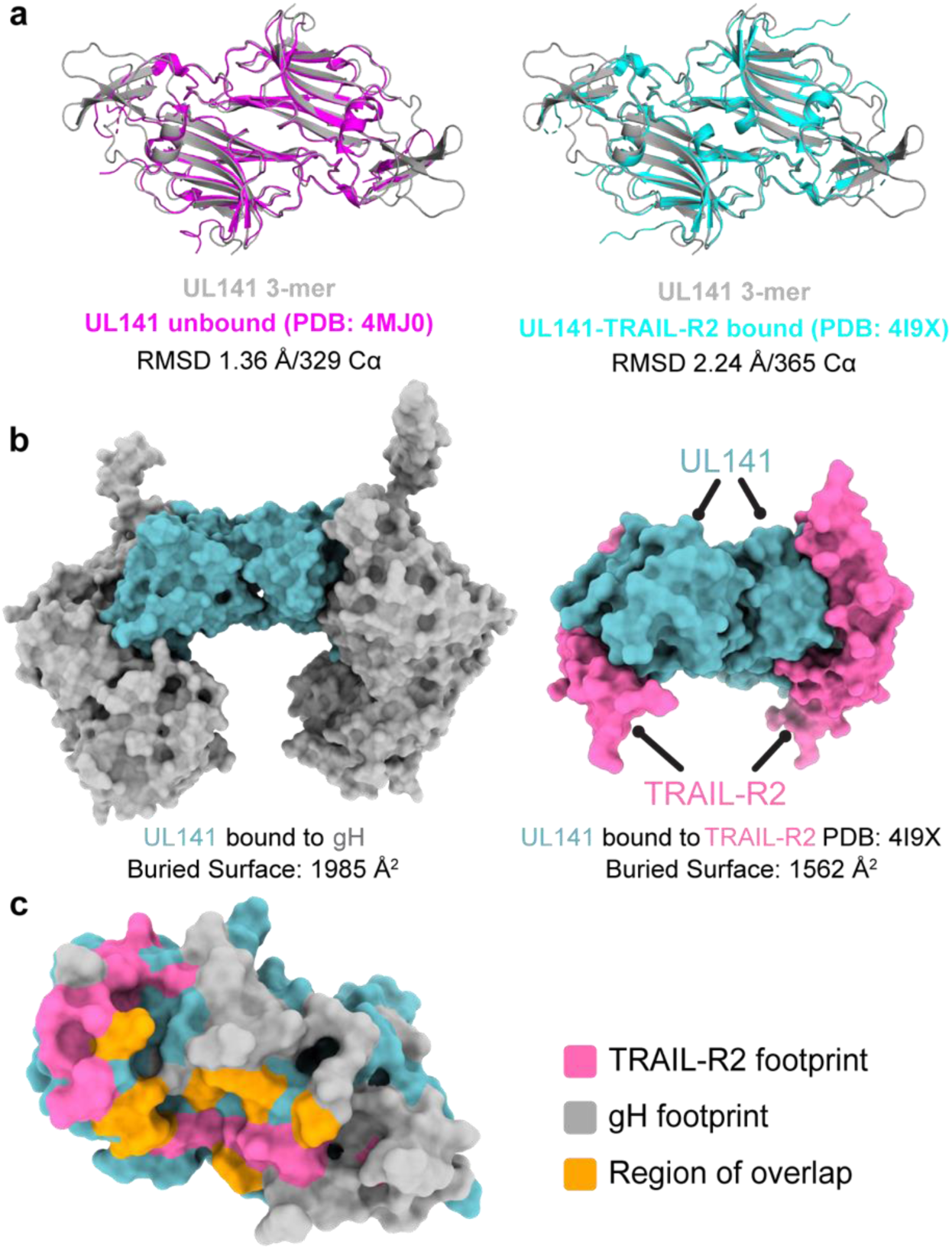
gH and TRAIL-R2 share a similar binding site on UL141. a, Structural comparison of UL141 from the 3-mer with unbound UL141 (left) and UL141 bound to TRAIL-R2 (right). The dimer structures were aligned as “dimers” using all Cα atoms in the respective PDB files, with the alignment quantified by the indicated r.m.s.d. values. **b,** Surface models of UL141 in the 3-mer (left) and UL141 bound to TRAIL-R2 (right) show that gH and TRAIL-R2 occupy similar binding sites on UL141. The calculated buried surface area for each interaction is indicated below. **c,** Surface model of a UL141 monomer with the TRAIL-R2 binding footprint highlighted in pink, the gH footprint in grey, and the region of overlap in orange. The calculated buried surface area of the overlapping region is indicated, accounting for approximately 25% of the TRAIL-R2 binding site.

## DISCUSSION

This study provides definitive evidence that there is a third functional complex on the HCMV envelope that facilitates entry and/or cell-to-cell spread in endothelial and epithelial cells, and is present in virions at higher levels than the trimer. This 3-mer viral surface complex is formed by interaction of the immunoevasin UL141 with gH and UL116. UL141 was previously only known to play an intracellular role in immune evasion. The studies presented here now illuminate that UL141 plays two critically distinct functions in the HCMV life cycle: (i) its previously known role in restricting expression of immune-activating molecules on the infected cell surface (Smith et al., 2013; Tomasec et al., 2005) and now (ii) its association with gH/UL116 in the virion envelope to enhance infectivity. Our results suggest that it is the UL141 component of the 3-mer that functions as a receptor binding moiety to enhance HCMV spread.

Cryo-EM of the 3-mer complex reveals a distinct and non-canonical organization. Although the 3-mer, trimer, and pentamer all share gH, the 3-mer alone lacks gL, and to our knowledge is the first example in any herpesvirus of a gH envelope complex where gL is absent. Instead, the 3-mer uses gH to scaffold its UL116 and UL141 binding partners, each on a different gH surface. Further, direct comparison of the HCMV 3-mer with the trimer or pentamer reveals a structural rearrangement of the gH DI to facilitate a stable interaction with UL116. This observed conformational change provides the structural basis for how UL116 can form a stable interaction with gH in the absence of gL, consistent with previous low-resolution studies (Caló et al., 2016).

The organization of UL141 within the 3-mer, where the binding of UL141 to gH obfuscates the expected binding site for TRAIL receptors, suggests that UL141 may not bind to these receptors when in its dimerized 3-mer form. However, our flow cytometry-based studies show that the recombinant 3-mer can bind to cell-expressed TRAIL-R1, –R2, –R4, and CD155 (**Fig. 14**). These results suggest the 3-mer may change conformation to facilitate receptor binding or separate from gH to allow receptor binding by UL141 alone. Uncovering this mechanism is a topic of current study. Plasticity in structure and partner interactions likely facilitates the ability of UL141 to play roles both inside the cell and out. Within the ER, UL141 interacts with human TRAIL-R2 to prevent this receptor and other immune molecules from reaching the infected cell surface. On the virion surface, UL141 instead interacts with gH and UL116 to form an abundant complex important for viral entry and cellular spread. Consequently, the many critical functions of UL141 suggest it may be a high-value vaccine target.

**Fig. 14.**
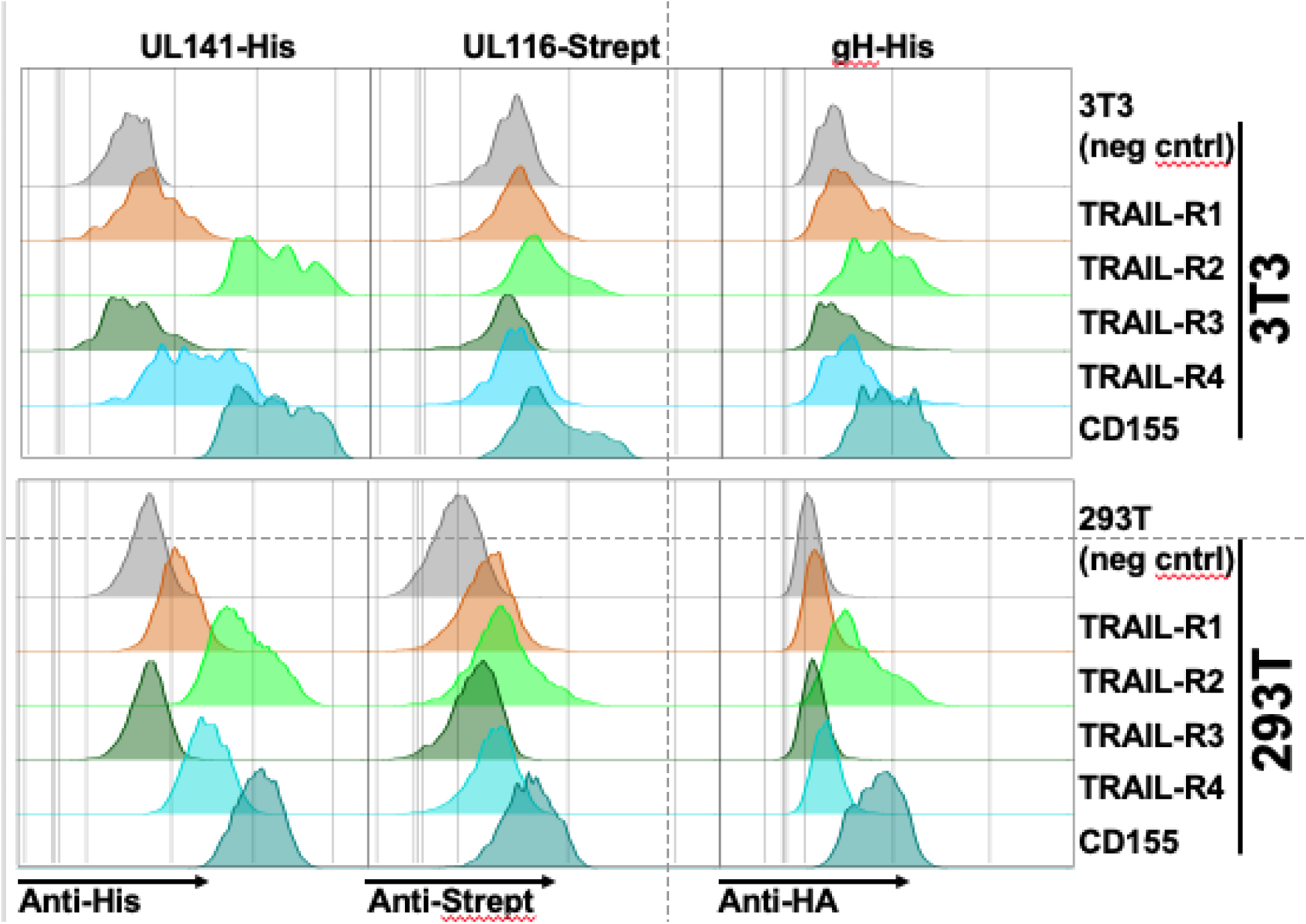
The 3-mer binds to UL141 interacting proteins. 3T3 or 293T cells were transfected with the four known human TRAIL receptors or CD155, using expression constructs where the receptor ectodomains were fused to a gpi-addition signal to facilitate cell-surface expression and avoid apoptosis mediated by overexpression of full-length TRAIL death receptors. Recombinant 3-mer protein engineered to express the indicated epitope tags on the individual subunits was then incubated with transfected cells (5μg/ml), and binding was detected using the indicated antibodies followed by flow-cytometry.

The structure of UL116 has not previously been solved at high-resolution. The cryoEM structure presented here reveals the C-terminal domain of UL116 to be wrapped about gH at the apex. Each UL116 monomer contains ∼31 kDa of polypeptide plus a full ∼120 kDa of carbohydrate (combined molecular weight of ∼150 kDa; 80% carbohydrate). The abundance of glycosylation and the apical positioning suggests that UL116 forms a type of glycan cap atop the 3-mer, perhaps to obscure this entry complex from antibody surveillance. Indeed, the abundance of glycosylation means that the N-terminal portion of the glycoprotein is mobile and disordered.

The independent scaffolding of UL116 and UL141, each by gH, invited the question of why UL116 is required for UL141 entry-promoting activity. We note that gH, without either gL or UL116, cannot mature beyond the ER (Caló et al., 2016; Kaye et al., 1992). Thus, we believe the role of UL116 in mediating UL141-dependent entry may lie in driving surface expression of the gH needed to scaffold UL141 and/or facilitate heterotrimerization of the 3-mer. The heavily glycosylated UL116 also forms a cap on top of the 3-mer and may promote recognition by lectins and/or shield the 3-mer from immune surveillance.

Cytomegalovirus lays a heavy burden on the global population, with 80% seroprevalence, lifelong infection, cognitive defects in children when their mothers are infected or re-infected in pregnancy, and a massive annual economic burden. In this work we reveal there is an additional glycoprotein complex on the viral surface that facilitates endothelial cell infection and is at least as abundant as the previously described trimer. CryoEM of the gH/UL116/UL141 complex reveals it to have a noncanonical organization. Design of useful vaccines against HCMV necessitates understanding of the different virion envelope protein complexes that mediate entry, so that immune responses are not easily escaped via entry by a different complex. The 3-mer provides the first new glycoprotein target for HCMV vaccine design in decades.

## METHODS

### Cell culture

Human telomerase immortalized human foreskin fibroblasts (HFFT) were prepared from primary HFF cells (ATCC, Cat # SCRC-1041) as previously described (Nguyen et al., 2018). ARPE-19 retinal pigment epithelial cells were purchased from ATCC (Cat # CRL-2302). HEK-293T cells were purchased from Genhunter Corp. (Nashville, TN). All cells were cultured in Dulbecco’s Modified Eagle’s Medium (DMEM, Corning) supplemented with 25 μg/mL gentamicin, 10 μg/mL ciprofloxacin, and 5% newborn calf serum (NCS, Sigma #N4637 or Gemini Bio GemCell). Pooled primary human umbilical vein endothelial cells (HUVEC) were purchased from Lonza (Cat # C2519A). HUVEC were grown and maintained in complete MCDB131 media supplemented with 10% fetal bovine serum (FBS, Sigma #F2442), 0.2% bovine brain extract, 10 mM HEPES, 60 ug/mL heparin, 2 mM glutamine, 25 μg/mL gentamicin, and 10 μg/mL ciprofloxacin. Heparin-free MCDB131 media was used for all infections of endothelial cells. Cell lines were routinely tested for mycoplasma using a PCR-based kit (Myco-Sniff-Valid™, MP Biomedicals).

### HCMV propagation and BAC reconstitution

Viruses were reconstituted by electroporation of HCMV bacterial artificial chromosomes (BACs) into HFFT, as described previously (Wang et al., 2013). Parental TB40/E (TB40-BAC4) (Sinzger et al., 2008), TR3 (Caposio et al., 2019), AD169 (AD169rv) (Hobom et al., 2000) and derivatives were amplified at low MOI on HFFT until extensive CPE was observed. Viruses were grown in complete DMEM. After 6 days of 100% CPE, cell-associated virus was released by freeze-thaw lysis and centrifuged at 3000*g* for 10 minutes to pellet cell debris. Supernatants were then ultracentrifuged in the SW 32 Ti rotor (24,000 rpm, 1.5 h, 4°C) through a 20% sorbitol cushion. The resulting virus pellets were resuspended in DMEM containing 5% NCS. Only passage 1 to passage 3 viral stocks were used for each experiment. Infectivity of virus stocks and samples were determined by the tissue culture infectious dose 50% (TCID_50_) assay. Briefly, serial dilutions of virus were used to infect HFFT in multiple wells of a 96-well plate. After 10 days, HFFT were stained for IE1 to score infected wells, and TCID_50_ values were calculated according to the Spearman-Kärber method.

For the purification of mature virions, viruses were propagated on HFFT cells and concentrated through a 20% sorbitol cushion after 6 days of 100% CPE. Virus pellets were resuspended in 2 mL sodium phosphate buffer, then layered on top of a positive density/negative viscosity glycerol/sodium tartrate gradient. The gradients were ultracentrifuged (SW 41 Ti) at 24,000 rpm for 1h at 10°C. A sterile needle was used to extract the third band containing infectious viral particles. The infectious particles were then pelleted through sodium phosphate buffer and resuspended in TX lysis buffer (0.1% Triton X-100, 50mM NaCl, 50mM Tris) for downstream immunoblotting.

### Generation of recombinant viruses

The TR3 BAC, a BAC clone of the clinical HCMV strain TR that has been fully restored to wild-type status and to which ganciclovir sensitivity was restored, was a generous gift of Dr. Klaus Frueh (Oregon Health and Science University, Beaverton, OR) (Caposio et al., 2019). We constructed a GFP-tagged TR3 by BAC mutagenesis. We first used *en passant* BAC recombineering to insert an excisable kanamycin resistance allele into the *GFP* expression cassette located between *US34* and *TRS1* in a BAC-cloned GFP expressing TB40-BAC4 derivative (Umashankar et al., 2011), now commonly called TB40E_5. This procedure entailed using primers EGFP_in_Kan_Fw and EGFP_in_Kan_Rv to PCR amplify a kanamycin resistance cassette that contains an *ISceI* recognition site at its 5’ end (*I-SceI-Kan)* (**Supplemental Data Table 2**). The DpnI digested PCR product was then electroporated into *Eschericia coli* GS1783 carrying the TB40E_5 BAC. After confirming that desired recombination event had occurred, a new PCR product containing the GFP cassette and portions of *US34* and the *US34/TRS1* intergenic region was prepared by PCR, using primers Us34CT_Fw and TRS1/Us34 Reg Rv (**Supplemental Data Table 2**). The PCR product was electroporated into E. coli GS1783 containing the TR3 BAC. Kanamycin resistant colonies were picked, grown up, confirmed and then resolved to remove the *I-SceI-Kan* cassette. Similar techniques were used to insert a FLAG (DYKDDDDK) epitope tag at the C-terminal cytoplasmic tail of UL141. Recombinant clones were confirmed by restriction fragment length polymorphism, PCR and Sanger DNA sequencing. Additionally, Illumina NextSeq550 sequencing at MiGS Center (Pittsburgh, PA) was carried out to exclude unexpected mutations within the entire viral genome.

### Plasmids

The TRAIL-R1:gpi and TRAIL-R2:gpi plasmids have been described (Bossen et al., 2006; Smith et al., 2013), and were gifts from Dr. Pascal Schneider. For structural studies, HCMV gH, UL116, and UL141 (strain TB40-BAC4 or Merlin) were subcloned into the phCMV mammalian expression vector containing an N-terminal Gaussia luciferase signal sequence (MGVKVLFALICIAVAEA) codon optimized for human expression. The gH gene comprised only the extracellular region (residues 30-709), the UL116 gene was truncated to lack the putative native signal sequence (residues 25-313), and the UL141 gene comprised only the extracellular region (residues 30-278). For protein purification purposes, UL116 was fused to a C-terminal enterokinase cleavage site and a Twin-StrepII-Tag and UL141 was fused to a C-terminal HRV-3C cleavage site and an 8X His-Tag. Plasmids were transformed into Stellar competent cells and isolated using a Plasmid Plus Midi kit (Qiagen).

### Protein Expression and Purification

The soluble ectodomain of the HCMV gH/UL116/UL141 trimer was purified in three steps. Plasmids encoding individual subunits were simultaneously co-transfected at an equimolar ratio into Freestyle 293-F cells (Thermo Fisher) using polyethyleneimine. At 5 days post-transfection, cultures were clarified by centrifugation at 4000 x g and supernatants were filtered through a 0.22 µm filter to remove cell debris prior to purification. Filtered supernatant was loaded on a 5 mL HisTrap-HP column (Cytiva), washed with 10 column volumes (CVs) of wash buffer (25 mM Tris pH 8.0, 150 mM NaCl, 10 mM imidazole), and eluted with 10 CVs of elution buffer (25 mM Tris pH 8.0, 150 mM NaCl, 250 mM imidazole). The eluate was applied to a 1 mL StrepTrap-HP column (Cytiva), washed with 5 CVs of Strep-wash buffer (25 mM Tris pH 8.0, 150 mM NaCl), and eluted with Strep-elution buffer (25 mM Tris pH 8.0, 150 mM NaCl, 5 mM d-desthiobiotin). The eluate was concentrated with an Amicon Ultra centrifugal filter device (30 kDa molecular weight cutoff [MWCO]) and loaded on a Superose 6 increase 10/300 column (Cytiva) equilibrated in size exclusion chromatography (SEC) buffer (25 mM Tris pH 8.0, 150 mM NaCl).

### Cryo-EM sample preparation and data acquisition

Purified gH/UL116/UL141 (3 uL) at a concentration of ∼0.8 mg/mL was deposited on holey carbon grids (C-Flat 2/1, copper 300 mesh, 40 nm carbon thickness). Excess liquid was blotted away for 6 seconds in a Vitrobot Mark IV operating at 4°C and 100% humidity before being plunge-frozen into liquid ethane cooled by liquid nitrogen. Movie stacks were collected using EPU (ThermoFisher) on a Titan Krios (ThermoFisher) operating at 300 keV with bioquantum energy filter equipped with a K3 direct electron detector at 1.1 Å/pixel and a total dose of 50.0 e/Å2.

### Cryo-EM data processing and model building

Recorded movies were corrected for frame motion using Cryosparc’s Patch-based motion correction and contrast transfer function (CTF) parameters were determined using Cryosparc’s Patch CTF program (Punjani et al., 2017). CTF-fitted images were filtered on the basis of the detected fit resolution better than 6 Å. Particles were picked using the TOPAZ neural network picker (Bepler et al., 2019) and cryoSPARC was used for two-dimensional (2D) classification, ab initio 3D reconstruction, 3D hetero-refinement, and non-uniform refinement. To further improve the map quality, particle coordinates were transferred to RELION 4 (Zivanov et al., 2018). In RELION, particles were sorted via 3D classification without alignment. To further improve the map around the gH/UL116 interface, particles were symmetry expanded and focused refinement was performed in cryoSPARC. Final half maps were used for local resolution estimation in cryoSPARC. Final reconstructions were sharpened with DeepEMhancer (Sanchez-Garcia et al., 2021) to aid in model building. A full description of the cryo-EM data processing workflows can be found in **Fig 6d**.

Refinement of each model was done through iterative rounds of manual model building using COOT 0.9.5 (Emsley et al., 2010) and ISOLDE (Croll, 2018), followed by refinement and validation performed in Phenix 1.20 (Liebschner et al., 2019). Model building at the gH/UL116 interface was aided by a protein model generated from AlphaFold2 (Evans et al., 2022; Jumper et al., 2021; Mirdita et al., 2022). Structural alignments and calculations of RMSD were carried out using the program PyMOL (http://www.pymol.org). The electrostatic surface representation was calculated in PyMOL using the APBS Electrostatics plugin (Baker et al., 2001; Dolinsky et al., 2007). ChimeraX-1.3 (Pettersen et al., 2021) was used to prepare figures of the structure. Full cryo-EM data collection and refinement statistics can be found in **Supplemental Data Table 3**.

### Absolute Infectivity Assays and Percent Infection Assays

HFFT or HUVEC were infected with the indicated viruses at MOI 1 or MOI 2, respectively. Supernatants were collected at 6 dpi and cleared of cell debris by centrifugation at 3,000 *g*. Prior to viral genome extractions, 200 µL of supernatants were treated with DNase I (NEB, catalog M0303L) for 30 minutes at 37°C to degrade free DNA (80% degradation in DMEM, 99% degradation in MCDB131). Viral genomes were then extracted in accordance with the manufacturer’s protocols for the PureLink^TM^ Viral RNA/DNA Mini Kit (Invitrogen, cat# 12280050). Viral genomes were quantified via qPCR using Luna Mastermix (NEB, cat# M3003L) and primers targeting the UL69 gene (110% efficiency) as described previously (Siddiquey et al., 2021). Standard curves were generated with 10-fold serial dilutions of parental TB40-BAC4 DNA to quantify viral genomes/mL. In parallel, supernatants were serially diluted to calculate TCID50/mL on HFFT, ARPE-19, and HUVEC. Infected cells were detected by staining for IE1 after 3 dpi. Ratios of TCID50/mL to genomes/mL measure the absolute infectivity of virions in TCID50/genome.

For percent infection assays, HFF-T and ARPE-19 were infected with 50 genome equivalents/cell while HUVEC were infected with 100 genome equivalents/cell with the indicated viruses. To exclude potential media effects, viruses were diluted in a 1:1 mixture of DMEM and heparin-free MCDB131 media. For microscopy, cells were stained with anti-IE1 (mouse monoclonal antibody 1B12) and counterstained with Hoechst 33342 at 1 dpi.

### Immunofluorescence microscopy

A detailed protocol for immunofluorescence microscopy has been described (Zhang et al., 2019). Fibroblasts were infected at MOI 1 with TB_UL141^FLAG^ virus, formalin fixed at 3 days post-infection (dpi) and blocked with human Fc block and 5% normal goat serum. Cells were imaged by confocal microscopy after staining with rabbit anti-UL141 antibody (1:1,000 dilution) or mouse anti-gB mAb clone 27-156 (1:800 dilution), which was a gift of William J. Britt (University of Alabama, Birmingham). Secondary antibodies used were Alexa Fluor 488 goat anti-mouse IgG (H+L) (Invitrogen, Cat# A11001) and Alexa Fluor 546 goat anti-rabbit IgG (H+L) (Invitrogen, Cat# A11010), each at 1:1000 dilution (**Supplemental Data Table 1**). Nuclei were counterstained using Hoechst 33342 (Thermo Fisher) prior to mounting coverslips with Prolong-Gold anti-fade reagent. Pictures were captured on a Nikon SIM-E and A1R confocal microscopy system with 100X/1.49 NA lens objective, z-stack scanning was applied.

### Statistics

Statistical details of experiments, including numbers of replicates and measures of precision, can be found in the figure legends, figures, Results, and Methods. All analyses were performed with GraphPad Prism version 10.2.3.

## Data Availability

All data needed to evaluate the conclusions in the paper are present in the paper and/or the Supplemental Data. Requests for resources and reagents should be directed to and will be fulfilled by the lead contacts, J.P.K., C.A.B., E.O.S. Structural models of the HCMV 3-mer have been deposited in the Protein Data Bank (www.rcsb.org) under accession numbers 9DIX (3-mer) and 9DIY (focused gH/UL116). Cryo-EM maps are deposited in the EM Database (https://www.emdataresource.org/) with the following ID: EMD-46920 (3-mer) and EMD-46921 (focused gH/UL116).

## Acknowledgements

We thank Ruben Diaz Avalos and the LJI CryoEM center for assistance with data collection. Equipment of the LJI cryoEM core was supported by NIH U19109762-S1, the GHR Foundation, and private donations. We thank the LSUHS Research Core Facility (RRID:SCR_024775) for assistance with confocal microscopy imaging and qPCR. We thank Jens von Einem (Universitätsklinikum Ulm, Ulm, Germany) and Gang Li (Johnson & Johnson, Rockville, Maryland) for expert advice on glycerol tartrate gradient purification of virions. This work was supported by NIH grants AI116851 to J.P.K., AI139749 and AI101423 to C.A.B., T32HL155022 to L.A.H., as well as institutional funds from the La Jolla Institute for Immunology (E.O.S. and C.A.B.), and from LSUHS in the form of a Chancellor’s Aim High award (J.P.K.) and an LSUHS Malcolm Feist Cardiovascular Fellowship (L.A.H.).

## Author contributions

J.P.K., C.A.B. and E.O.S. conceived the idea, developed and supervised the project. M.N.A.S. constructed recombinant HCMVs and carried out immunoprecipitation experiments. L.A.H. carried out absolute infectivity and percent infection assays, glycerol/sodium-tartrate purification of virions, Western blotting, confocal microscopy imaging and analysis, viral growth curves, and plaque size measurements. M.N. constructed new expression plasmids, purified proteins, carried out cryo-EM studies, and solved the 3-mer structure. J.Y. assisted with cloning and 3-mer purification. S.B. and K.Y. performed various binding/biochemical studies with the purified 3-mer.

## FIGURE LEGENDS

**Supplemental Data Fig. S1.** UL141 localizes at the cVAC. Immunofluorescent staining of fibroblasts infected with TB40 viruses that are UL141-null (TB40^141-^) or express FLAG-tagged UL141 (TB40^141F^) at 3 dpi (MOI 1 TCID50). Cells were stained with anti-UL141 (magenta) and anti-gB (green) or anti-FLAG (magenta) and calnexin (CNX, green).

**Supplemental Data Fig. S2. UL141 promotes viral spread independently of the pentamer complex. a**, Plaque sizes recorded for each biological replicate in AD169 and AD169^141^ infected ARPE-19 cells at 10 dpi. After log_10_ transformation, data fit a Gaussian distribution and are used to calculate statistical significance via 2-way ANOVA. **b**, QQ plots displaying the lognormality of raw plaque size data.

