## Supplemental Data Table 1 for "A noncanonical glycoprotein H complex enhances cytomegalovirus entry"

**Extended Data Table 1**. Primary antibodies and working dilutions used for each application.

| Antibody Name or Target | Animal Source | Mono-/ Polyclonal | CloneID or Rabbit ID | Vendor or Source | Cat. No. | Dilutions for Applications | | | |
|---|---|---|---|---|---|---|---|---|---|
| | | | | | | *IP* | *WB* | *IF* | *FC* |
| DYKDDDDK (aka FLAG) tag | Rb | monoclonal | D6W5B | Cell Signaling Technology | 14793 | 1:50 | 1:1000 | | |
| FLAG tag | Ms | monoclonal | M2 | Sigma Aldrich | F1804 | | 1:1000 | 1:1000 | |
| ECS (DYKDDDDK) Tag | Rb | Monoclonal | BLRE00G | Bethyl | A191-100 | 1:100 | | | |
| Myc-tag | Ms | monoclonal | 9B11 | Cell Signaling Technology | 2276S | 1:250 | 1:1000 | 1:8000 | |
| His-tag | Ms | monoclonal | J099B12 | BioLegend | 652502 | | 1:1000 | | |
| Strep-tag | Ms | monoclonal | GT661 | GeneTex | GTX628900 | | 1:1000 | | |
| HA-tag | Ms | monoclonal | J095G46 | Biolegend | 362605 | | | | 5µg/ml |
| IE1 | Ms | monoclonal | 1B12 | Thomas Shenk, Princeton University | n/a | | 1:200 | 1:200 | |
| UL141 (residues 308-338) | Rb | polyclonal anti-peptide | 14501 | Pacific Immunology | n/a | | 1:1000 | 1:1000 | |
| UL141 | Ms | monoclonal | M550.2 | Dr. Wilkinson, Cardiff Univ | n/a | | 1:1000 | | 5µg/ml |
| gB | Ms | monoclonal | 27-156 | William J. Britt, University of Alabama, Birmingham | n/a | | 1:1000 | 1:800 | |
| gL (residues 258-278) | Rb | polyclonal anti-peptide | 11684 | Pacific Immunology | n/a | | 1:1000 | | |
| MCP | Ms | monoclonal | 28-4 | William J. Britt, University of Alabama, Birmingham | n/a | | 1:400 | | |
| UL148 (residues 263-285) | Rb | polyclonal anti-peptide | 9220 | Pacific Immunology | n/a | | 1:1000 | | |
| gH | Ms | monoclonal | AP86 | William J. Britt, University of Alabama, Birmingham | n/a | | 1:1000 | 1:800 | |
| gH | Ms | monoclonal | 11-1-1 | Dr. Britt, AL Birmingham | n/a | | | | 1:200 |
| Calnexin | Rb | monoclonal | C5C9 | Cell Signaling Technology | 2679S | | 1:1000 | 1:1000 | |

Key: Rb: Rabbit; Ms: Mouse
