## Supplemental Data Table 2 for "A noncanonical glycoprotein H complex enhances cytomegalovirus entry"

**Extended Data Table 2. Oligonucleotide Primers for HCMV BAC recombineering procedures.**

| Primer Name | Purpose | PCR template | BAC targeted for modification | Sequence, 5'-3' |
|---|---|---|---|---|
| UL141 repair Kan Fw | Repair UL141 frameshift in HCMV strain TB40/E | I-SceI-AphAI (KanR) sequence (PMID: 16526409) | TB40-BAC4 (PMID: 18198366) or derivatives therof | GTCGCCGGCGCCGGTGTTGG TCGCCGAGGGAGAGCAAGTTA CCATCCCCTGCACGGTCATAG GGATAACAGGGTAATCGATTT |
| UL141 repair Kan Rv | | | | CCATGGGCCAGGAGTGTGTCA TGACCGTGCAGGGGATGGTAA CTTGCTCTCCCTCGGCGAGCC AGTGTTACAACCAATTAACC |
| UL141_FLAG_Kan_Fw | Add FLAG epitope tag to the C-terminus of UL141 | I-SceI-AphAI (KanR) sequence (PMID: 16526409) | TB40-BAC4 (PMID: 18198366), TR3 (PMID: 31848362) or derivatives thereof | GACTTACGATAGTTACCCCGGT GTTAAAAAGATGAAGAGGGAC TACAAGGATGACGACGATAAGT GAGAACATAGGGATAACAGGG TAATCGATTT |
| UL141_FLAG_Kan_Rv | | | | TTTTTTAACATGTTATTTTTTTATTT TATGCGTGTTCTCACTTATCGTCG TCATCCTTGTAGTCCCTCTTCAGC CAGTGTTACAACCAATTAACC |
| Xfer_UL141_FLAG_Fw | Replace UL131-UL128 with UL141-FLAG CDS | TR3-UL141-FLAG-Kan-integrate (intermediate in BAC recombineering) | AD169rv (Hobom et al., 2006, PMID: 10933677) | ATGATGTCTCATAATAAAGCTTTCT TTCTCAGTCTGCAACAGCGTCTC TGCGAAAAAGG |
| Xfer_UL141_FLAG_Rv | | | | CGACAGAAATCTCAAAACGCGTAT TTCGGACAAACACACACCAACAC GCCCATTCAATCC |
| EGFP_in_Kan_Fw | To produce an I-SceI-AphAI-disrupted eGFP locus in a GFP+ TB40-BAC4 derivative to enable transfer of the eGFP locus to the TR3 BAC | I-SceI-AphAI (KanR) sequence (PMID: 16526409) | TB40E_5: a TB40-BAC-4 derivative expressing eGFP (PMID: 22241980) | GGCAACTACAAGACCCGCGCCG AGGTGAAGTTCGAGGGCGACAC CCTGGTGAACCGCATCTAGGGAT AACAGGGTAATCGATTT |
| EGFP_in_Kan_Rv | | | | AAGTCGATGCCCTTCAGCTCGAT GCGGTTCACCAGGGTGTCGCCC TCGAACTTCACCTCGGCCAGTGT TACAACCAATTAACC |
| Us34CT_Fw | Transplant the AphAI-disrupted eGFP expression cassette from TB40-BAC4 GFP to TR3 | TB40E_5_eGFP-I-SceI-AphAI, a TB40-BAC-4 derivative expressing eGFP (PMID: 22241980) in which the eGFP locus is disrupted by an I-SceI-AphAI sequence that can later be removed via en passant BAC mutagenesis | TR3 (PMID: 31848362) | TGTTCCTCCTCTTGAACCGC |
| TRS1/Us34 | | | | CTCCGCATCCCACCATTCTT |
