## Supplemental Data Table 3 for "A noncanonical glycoprotein H complex enhances cytomegalovirus entry"

**Extended Data Table 3. Cryo-EM data collection, refinement, and validation.**

| | gH/UL116/UL141 3-mer | gH/UL116 Local |
|---|---|---|
| ***Data Collection*** | | |
| Microscope | Titan Krios G3 | |
| Voltage (keV) | 300 | |
| Detector | Gatan K3 | |
| Energy Filter | GIF Bioquantum | |
| Magnification (nominal/calibrated) | 130,000x | |
| Data acquisition software | EPU | |
| Exposure (s) | 2.5 | |
| Total electron exposure ($e^{-}/Å^{2}$) | ~50 | |
| Number of frames per micrograph | 50 | |
| Pixel size (raw/final) (Å) | 0.66/1.32 | |
| Defocus range (μm) | -0.6 to -2.5 | |
| Micrographs collected (no.) | 14,075 | |
| ***Reconstruction*** | | |
| Image processing package | CryoSPARC/Relion | CryoSPARC |
| Number of particles (unique/symmetry expanded) | 119,525 / - | 56,333 / 112,666 |
| Symmetry imposed | C2 | C1 from symmetry expanded C2 |
| Resolution (Å) | | |
| FSC 0.143 (unmasked/masked) | 4.2/3.5 | 5.4/6.3 |
| Local resolution range (min/median/max) | 2.9/4.5/35.1 | 3.7/5.1/7.4 |
| ***Model composition*** | | |
| Non-hydrogen Atoms | 29,158 | 9,037 |
| Protein Residues | 1802 | 546 |
| Ligands | BMA: 2, NAG: 16 | BMA: 1, NAG: 13 |
| ***Model vs. Data*** | | |
| CC (mask) | 0.76 | 0.58 |
| CC (box) | 0.82 | 0.72 |
| CC (peaks) | 0.69 | 0.38 |
| CC (volume) | 0.76 | 0.60 |
| Mean CC for Ligands | 0.77 | 0.80 |
| ***Model Refinement*** | | |
| Model-map FSC 0.5 (unmasked/masked) | 3.97/3.82 | 7.45/7.34 |
| Model-map FSC 0.143 (unmasked/masked) | 3.55/3.49 | 5.84/5.73 |

| | | |
|---|---|---|
| RMSD from ideal geometry | | |
| Bond length (Å) | 0.004 | 0.003 |
| Bond angles (°) | 0.611 | 0.780 |
| Ramachadran plot | | |
| Outliers (%) | 0.00 | 0.00 |
| Allowed (%) | 4.82 | 7.25 |
| Favored (%) | 95.18 | 92.75 |
| MolProbity score | 1.71 | 2.08 |
| Rotamer Outliers (%) | 0.56 | 0.21 |
| Clashscore (all atoms) | 6.77 | 12.88 |
| C-beta Outliers (%) | 0.00 | 0.00 |
| CaBLAM Outliers (%) | 2.20 | 4.58 |
| Refinement package | Phenix | Phenix |
