## Supplemental Data Fig. S1 for "A noncanonical glycoprotein H complex enhances cytomegalovirus entry"

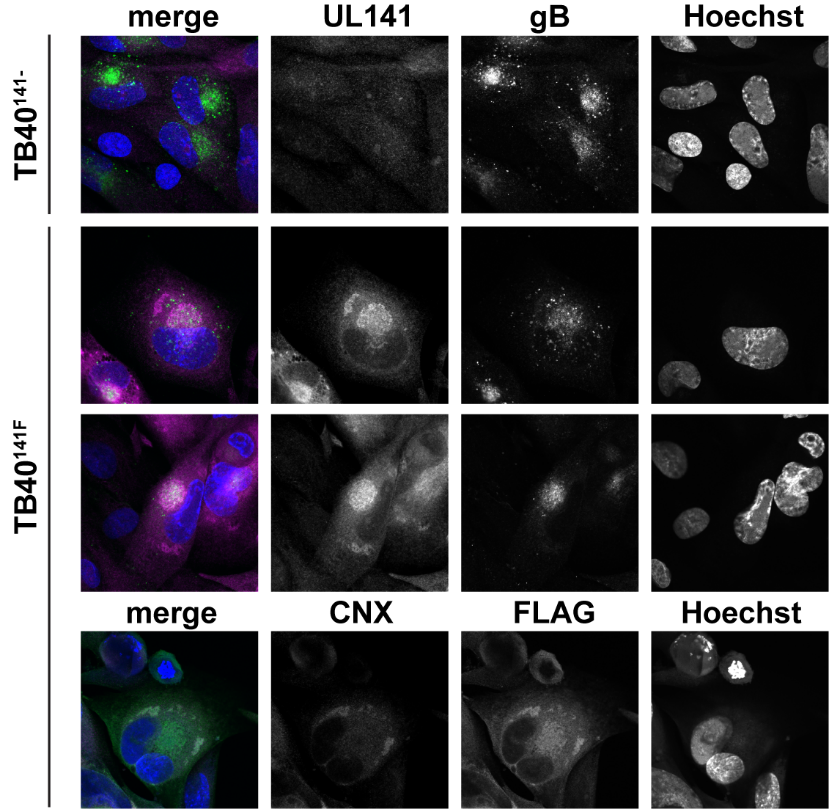

**Supplemental Data Fig. S1.** UL141 localizes at the cVAC. Immunofluorescent staining of fibroblasts infected with TB40 viruses that are UL141-null (TB40^141-^) or express FLAG-tagged UL141 (TB40^141F^) at 3 dpi (MOI 1 TCID50). Cells were stained with anti-UL141 (magenta) and anti-gB (green) or anti-FLAG (magenta) and calnexin (CNX, green).
