## Supplemental Data Fig. S2 for "A noncanonical glycoprotein H complex enhances cytomegalovirus entry"

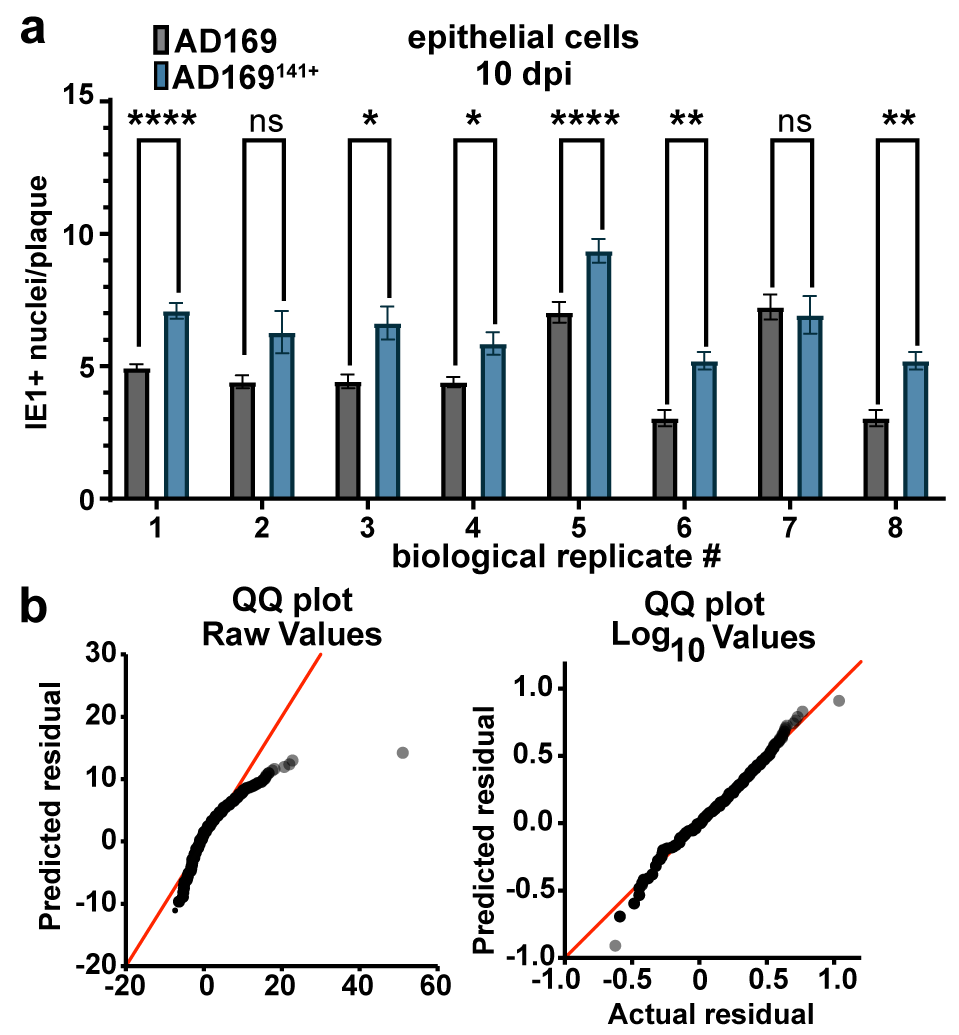

**Supplemental Fig. S2. a**, Plaque sizes recorded for each biological replicate in AD169 and AD169^141^ infected ARPE-19 cells at 10 dpi. After log_10_ transformation, data fit a Gaussian distribution and are used to calculate statistical significance via 2-way ANOVA. **b**, QQ plots displaying the lognormality of raw plaque size data.
